# Interferometric Ultra-High Resolution 3D Imaging through Brain Sections

**DOI:** 10.1101/2025.02.03.636258

**Authors:** Hao-Cheng Gao, Fan Xu, Xi Cheng, Cheng Bi, Yue Zheng, Yilun Li, Tailong Chen, Yumian Li, Alexander A. Chubykin, Fang Huang

## Abstract

Single-molecule super-resolution microscopy allows pin-pointing individual molecular positions in cells with nanometer precision. However, achieving molecular resolution through tissues is often difficult because of optical scattering and aberrations. We introduced 4Pi single-molecule nanoscopy for brain with *in-situ* point spread function retrieval through opaque tissue (4Pi-BRAINSPOT), integrating 4Pi single-molecule switching nanoscopy with dynamic *in-situ* coherent PSF modeling, single-molecule compatible tissue clearing, light-sheet illumination, and a novel quantitative analysis pipeline utilizing the highly accurate 3D molecular coordinates. This approach enables the quantification of protein distribution with sub-15-nm resolution in all three dimensions in complex tissue specimens. We demonstrated 4Pi-BRAINSPOT’s capacities in revealing the molecular arrangements in various sub-cellular organelles and resolved the membrane morphology of individual dendritic spines through 50-µm transgenic mouse brain slices. This ultra-high-resolution approach allows us to decipher nanoscale organelle architecture and molecular distribution in both isolated cells and native tissue environments with precision down to a few nanometers.

## Introduction

The brain, a network comprising billions of interconnected neurons, forms the basis for cognitive development ^1^. A neural circuit, the basic functional unit of the brain, consists of neurons interconnected by synapses where information flows from one neuron to another. The synapses facilitate the transmission of neurotransmitters from presynaptic terminals to postsynaptic spines^2^. Understanding the nano-anatomy of dendritic spines is crucial for comprehending brain development and higher functions ^3^. These spines, which house the postsynaptic components of most excitatory synapses, change in size throughout life, reflecting synaptic strength and plasticity during development and aging. Both the morphology of spine heads and the spine necks play critical roles in synapse transmission by regulating chemical and electrical compartmentalization ^4–8^. However, studying these protein complexes, organized at a nanometer scale of 15-60 nm inside thick tissues, poses challenges for conventional microscopy systems ^9^.

Due to the wave nature of light, the resolution of conventional fluorescence microscopy methods is limited to sub-micrometer level ^10–13^, inadequate to visualize synaptic ultrastructure and molecular distributions. Recent advancements in super-resolution microscopy techniques, such as stimulated emission depletion (STED) microscopy, single-molecule switching nanoscopy (SMSN), and structured illumination microscopy (SIM), have enhanced resolution by nearly tenfold ^14–18^. These techniques have empowered researchers to probe protein organization within presynaptic boutons and postsynaptic densities, primarily in isolated neuronal cultures ^19–25^.

Images of isolated neuron cultures have shed light on the spatial arrangements of proteins, synaptic interactions, and neuronal communication, deepening our understanding of neural circuitry ^9,24–26^. However, the extraction of individual neurons from their native tissue context compromises the structural and functional integrity of neural connections, hindering the investigation of synaptic regulation during information processing and memory formation. To uncover the molecular mechanisms of the circuit function in the native or disease contexts, direct examination inside brain tissue is imperative. Yet, resolving the molecular components within brain tissues is challenging due to the complex optical properties of intra-and-extra cellular constituents. Moreover, conventional 3D super-resolution microscopy techniques exhibit pronounced disparities between axial and lateral resolutions, especially through deep tissues, and are limited to a 3D resolution of ∼85-120 nm in brain sections of 50 μm or thicker ^27–34^. In contrast, to unravel the molecular intricacies of the brain, an optical imaging platform must attain and sustain an isotropic 3D resolution of sub-15 nm in unperturbed tissue specimens while minimizing distortions and artifacts.

Here, we have developed the 4Pi single-molecule nanoscopy for brain with *in-situ* point spread function retrieval through opaque tissue (4Pi-BRAINSPOT). This approach allows an *in-situ* localization precision of 6 nm laterally and 2.5 nm axially through 50-μm brain slices by synergizing the isotropic 3D resolution capabilities of 4Pi single-molecule switching nanoscopy (4Pi-SMSN) with a novel *in-situ* coherent PSF retrieval approach, SMSN-compatible tissue clearing, and highly inclined light-sheet illumination. We demonstrated that 4Pi-BRAINSPOT enables 3D nanoarchitecture visualization through thick brain tissues, revealing the molecular distribution and ultrastructure of dendritic spines while preserving circuit integrity unattainable in cultured neurons. We further developed a novel analysis pipeline for quantifying spine morphology utilizing the highly accurate molecular locations obtained by 4Pi-BRAINSPOT. We expect that 4Pi-BRAINSPOT will enable molecular-level investigation of synaptic regulation within neurobiological processes and thus enhancing our understanding of the mechanisms underlying various neurodevelopmental disorders.

## Result

### Design of 4Pi-BRAINSPOT

4Pi-BRAINSPOT (Figure 1A) is developed based on 4Pi-SMSN, which coherently detects the emitted photons from single emitters through two conjugated objectives ^35–39^ achieving sub-10 nm localization precision in all three dimensions. However, 4Pi-SMSN through tissues remains difficult. Biological tissues have complex optical properties, causing scattering and aberrations and thus deteriorating the interferometric emission pattern ^40,41^. Additionally, temperature-induced changes in the coherent cavity and persistent objective misalignments also affect the achievable interferometric contrast and emission pattern of single molecules.

**Figure 1.**
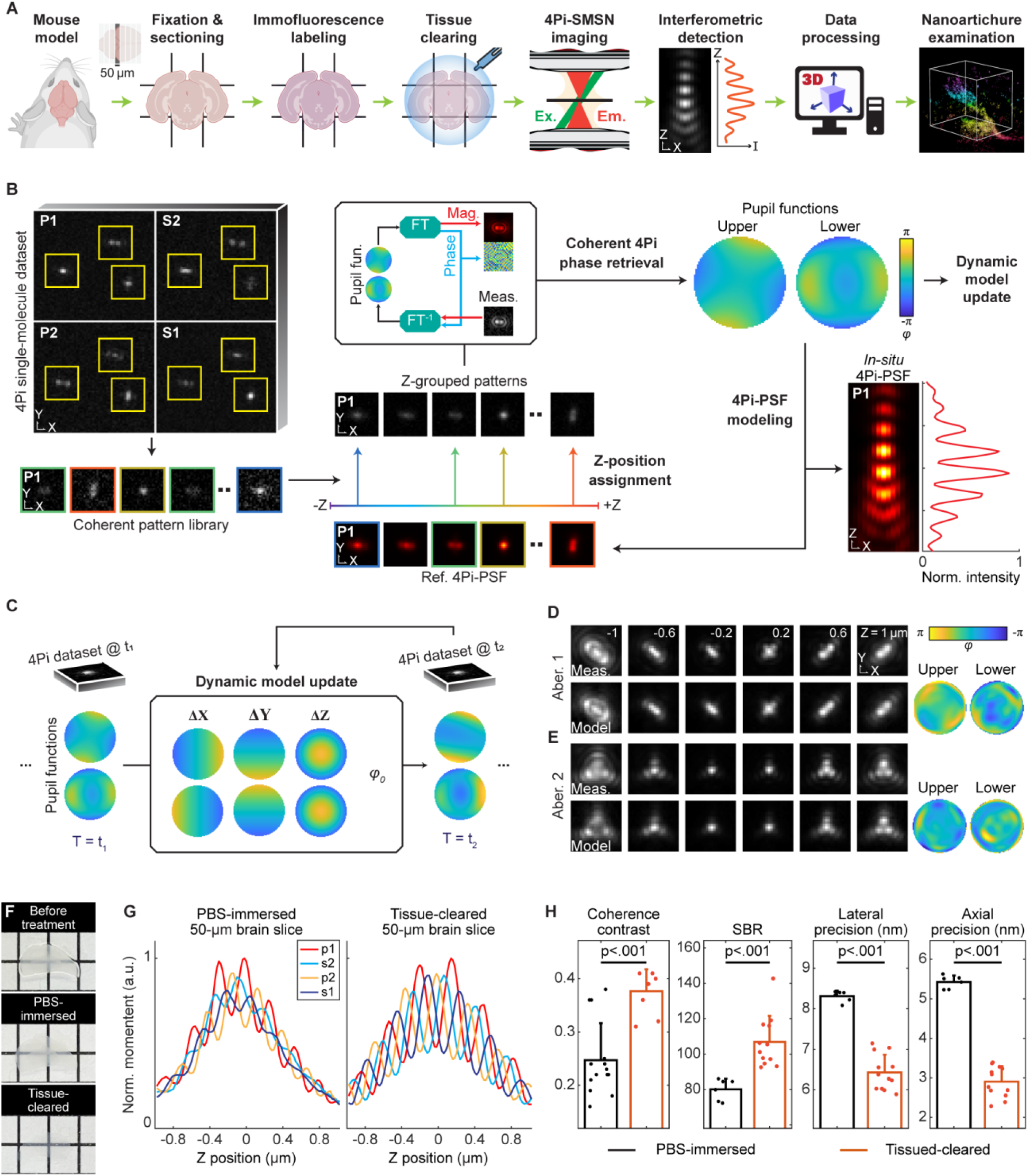
Overview of 4Pi-BRAINSPOT concepts and demonstrations. (**A**) Imaging workflow of 4Pi-BRAINSPOT. To image opaque brain tissues, we first excised, fixed, and sectioned the tissues into 50-μm coronal sections. These sections underwent immunofluorescence-labeling targeting the protein of interest, followed by tissue clearing, and were mounted with thiol-based imaging buffer. Imaging is conducted with 4Pi-SMSN, utilizing a 54°-inclined light-sheet illumination. The obtained single-molecule datasets were processed with the algorithm detailed in (**B**), which extracted dynamic *in-situ* retrieval of the 4Pi PSF model and facilitated the 3D localization of single molecules for detailed visualization and quantification. (**B**) Data processing workflow of *in-situ* PSF modeling in 4Pi-BRAINSPOT. The algorithm segmented individual 4Pi emission patterns from single-molecule datasets to create a library. These segments were assigned to specific axial groups according to their similarity to the reference PSFs generated from initial coherent pupil functions. The categorized patterns were then aligned and averaged within each group and underwent the coherent 4Pi phase retrieval algorithm (see also Figure **S2**), updating the individual pupil functions and reference PSFs. Through sequential iterations, the model was refined to represent *in-situ* interferometric emission patterns. The model was fed into the dynamic model update in (**C**), capturing time-varying response functions. (**C**) Data processing workflow of dynamic *in-situ* model update in 4Pi-BRAINSPOT. We generated a series of reference PSFs for each time window based on coherent pupil functions from the previous window, with adjustments of 3D objective misalignments (ΔX, ΔY, and ΔZ) and cavity phase shifts (φ_0_). The algorithm iteratively refined the time-dependent parameters to capture the alterations in single-molecule patterns according to the similarity between the PSF model and segmented emission patterns in the current time window. See also Figure **S3**. (**D**&**E**) Interferometric pattern comparison between fluorescent bead measurements (top row) and 4Pi-BRAINSPOT PSF models (bottom row) in channel p1 with two different aberrations at axial positions ranging from ±1 µm. Retrieved coherent pupil functions are shown on the right. See also Figures **S1A**-**G**. (**F**) Tissue-clearing results between mouse brain slices immersed in PBS and tissue-cleared mouse brain slices. Box size: 2.5 mm. (**G**) Gaussian-weighted moment operators of 4Pi interference of fluorescent bead measurement with PBS-immersed (left) and tissue-cleared (right) mouse brain slices. (**H**) Comparison of coherence contrast, signal-to-background ratio (SBR), and localization precisions between PBS-immersed (black) and tissue-cleared (orange) mouse brain slices. Bar height is the mean and error bar is the standard deviation. Statistical significance: t-test. See also Figure **S6**.

To mitigate tissue-induced aberration and temporal fluctuations in the interferometric system, we developed the 4Pi *in-situ* point spread function (PSF) retrieval algorithm, which enables direct retrieval of an aberrated, time-dependent *in-situ* model of the single-molecule emission patterns from the 4Pi-SMSN dataset captured in brain specimens (Figures 1B&C; Method 3.1). By deriving the model *in situ*, it reflects the distortions, blurring, and partial coherence encountered during detection, and thereby minimizes disparities between the data and the model. This approach provides accurate molecular localizations in the presence of tissue-induced aberrations.

The 4Pi *in-situ* PSF retrieval algorithm begins by segmenting multichannel 4Pi emission patterns to create a library of detected emission patterns (Method 3.2). These segments, from stochastically blinking events at varying axial positions, serve as random samples of the underlying *in-situ* interferometric (4Pi) PSF that we aim to retrieve. To estimate axial positions of patterns in the library, reference PSFs are generated from coherent pupil functions, depicting the interferometric amplitude and phase of wavefield superposition from the pupils of the two opposing objectives (Method 3.3) ^27,42,43^. The detected emission patterns are then assigned to specific axial groups according to their similarity to the reference PSFs. The categorized patterns are then aligned and averaged within each group to generate experimental PSF observations. To retrieve the individual pupil functions, we developed a coherent 4Pi phase retrieval algorithm based on the Gerchberg–Saxton algorithm. This allows us to retrieve individual pupil functions for each objective from their coherent superpositions and update the reference 4Pi-PSFs for the subsequent iteration (Figure S3; Method 3.4). Through sequential iterations, the model is refined to capture detailed coherent aberrations from both instrumental imperfections and biological specimens, representing *in-situ* interferometric emission patterns (Figures 1D&E) for accurate and precise single-molecule localization inside tissue specimens (Figures S4C; Method 3.7).

To address temporal aberrations induced by the system and specimens, commonly encountered due to the extended acquisition time of 4Pi-SMSN, 4Pi-BRAINSPOT employs a dynamic *in-situ* model update to mitigate drift-induced disparities between the acquired 4Pi data and the 4Pi-PSF model. In brief, 4Pi-BRAINSPOT detects and iteratively updates the extracted coherent pupil functions to capture the temporal alterations of single-molecule patterns caused by time-dependent objective misalignments and interferometric cavity shifts (Figure S4A&B; Method 3.5&6). Compared to traditional fiducial-bead-based *in-vitro* approaches, these dynamic refinements result in a time-varying response function of the 4Pi-SMSN unique to each time point.

4Pi-BRAINSPOT further incorporates an SMSN-compatible tissue-clearing method to mitigate aberration and scattering within opaque specimens, thereby preventing the permanent loss of Fisher information ^41^. We found that a combination of fast optical clearing method (FOCM; Method 1.8) ^44^, facilitated by protein denaturation and tissue hyperhydration, with a thiol-based imaging buffer significantly reduces tissue heterogeneity while preserving the photo-switching behavior of fluorophores critical for SMSN experiments (Figure 1F). Comparative results with and without FOCM inside 50-μm thick samples (Figures 1G&H) indicate a 1.5-fold increase in coherent fringe contrast (from 0.25±0.07, mean ± SD, n=14 beads, to 0.38±0.04, n=8 beads) and a 1.3-fold increase in signal-to-background ratio (SBR, from 80.1±6.0, n=6 datasets, to 106.9±14.7, n=12 datasets). Furthermore, by introducing highly inclined and laminated optical sheet (HILO) as excitation ^45^, 4Pi-BRAINSPOT further enables a 60% reduction in out-of-focus background across 50-μm thick brain sections (Method 4.3). These improvements collectively led to a localization precision of 6.4±0.4 nm laterally and 2.9±0.4 nm axially as quantified through tissue specimens (n=12 datasets; Figure S2).

### Characterization of 4Pi-BRAINSPOT performance

We first tested the accuracy and precision of *in-situ* PSF modeling by retrieving known distortions from simulated 4Pi emission patterns with random axial positions in a range of ±800 nm (Figure 2A). The simulated aberrations included 11 Zernike modes with random amplitudes between ±1 λ/2π, with vertical astigmatism as a known prior breaking the axial ambiguity (Method 2.6). Our methodology successfully retrieved the intricate fringes of the interferometric patterns (Figure 2B-D), achieving a normalized cross-correlation similarity (NCC) of 0.98 ± 0.05 (n=30 trials; Method 3.3) comparing to the ground truth, consistently outperforming conventional *in-vitro* approaches (NCC of 0.75 ± 0.20, n=60 trials). Furthermore, we assessed the algorithm’s ability to retrieve lateral and axial objective misalignments (ΔXY & ΔZ) and dynamic system shifts as cavity phase differences (φ0) (Figures 2E-G). Our approach allowed us to retrieve these commonly encountered interferometric system drifts at lateral and axial directions with standard deviations of 4.4 nm and 1.4 nm (n=60 trials) and captured the cavity phase difference consistently and accurately with a standard deviation of 12.7 mrad (n=60 trials). The algorithm also accurately captured the cavity phase with deviated initial guesses, indicating minimal influence of initial guess in obtaining an accurate PSF model (Figure 2H; Method 2.6).

**Figure 2.**
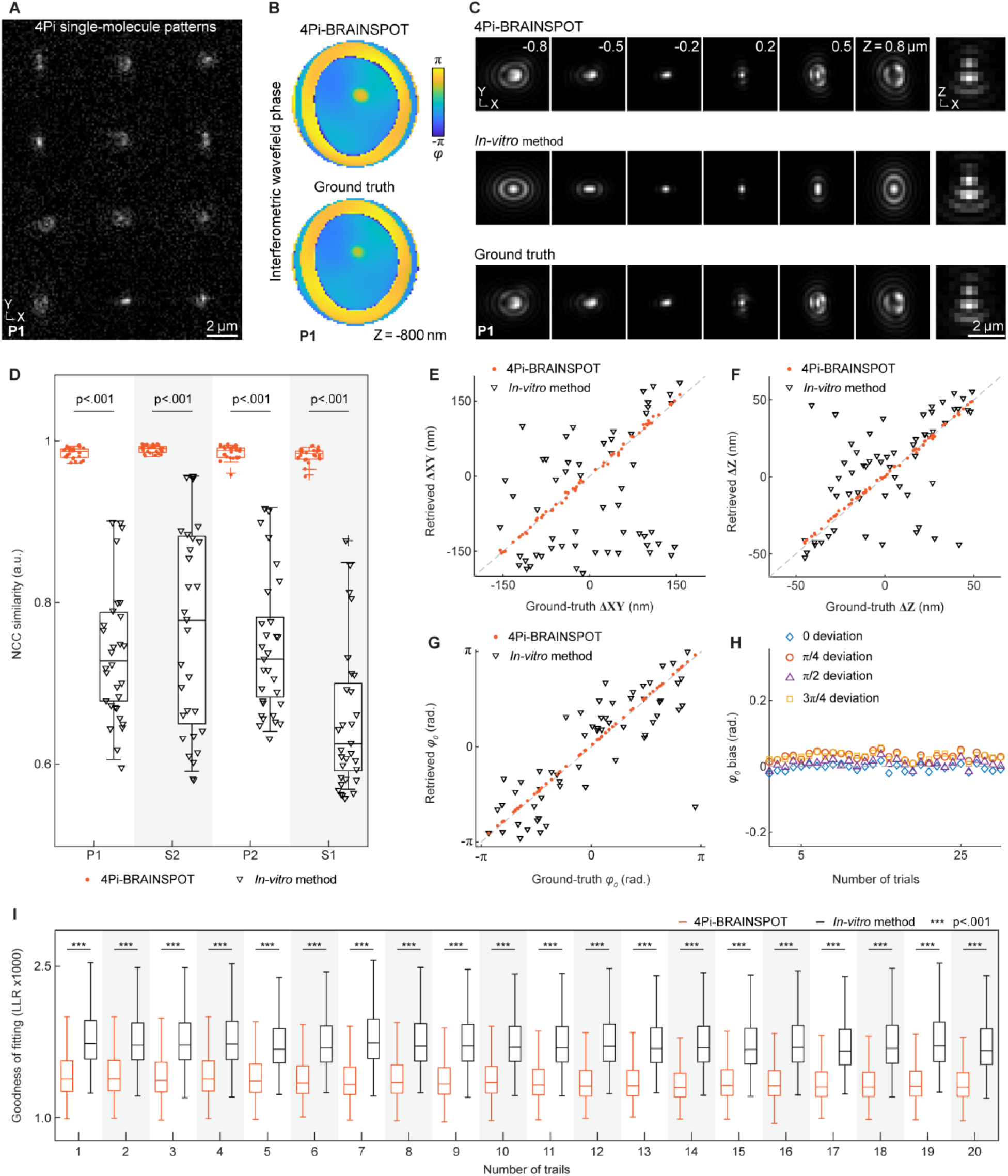
Performance quantification of 4Pi-BRAINSPOT. (**A**) Simulated 4Pi single-molecule emission patterns in channel p1 with known aberrations at random axial positions ranging from ±800 nm. (**B**) Comparison of interferometric wavefield superpositions at the axial position of -800 nm in (**A**) retrieved using 4Pi-BRAINSPOT (top) and the ground truth (bottom). (**C**) Horizontal and vertical cross-sections of PSFs of the 4Pi-BRAINSPOT (top row), *in-vitro* method (middle row), and ground truth (bottom row). (**D**) NCC similarity between ground truth PSFs and modeled PSFs using 4Pi-BRAINSPOT (orange) and *in-vitro* method (black). Boxes are upper and lower quartiles, central lines are median, and error bars are furthest data points excluding outliers (n=30 trials for each). Statistical significance: t-test. (**E**) Scatter plot of ground-truth lateral objective misalignment versus retrieved objective misalignment in simulations using dynamic *in-situ* model update with coherent pupil functions retrieved using 4Pi-BRAINSPOT (orange) and *in-vitro* method (black) (n=60 trials for each). (**F**) Scatter plot of ground-truth axial objective misalignment versus retrieved objective misalignment in simulations using dynamic *in-situ* model update with coherent pupil functions retrieved using 4Pi-BRAINSPOT (orange) and *in-vitro* method (black) (n=60 trials for each). (**G**) Scatter plot of ground-truth phase cavity versus retrieved objective misalignment in simulations using dynamic *in-situ* model update with coherent pupil functions retrieved using 4Pi-BRAINSPOT (orange) and *in-vitro* method (black) (n=60 trials for each). (**H**) Bias of cavity phase estimation with deviated initial guess of coherent pupils of 0, π/4, π/2, and 3π/4 (n=30 trials for each). (**I**) Goodness of fitting between the obtained emission patterns and modeled PSFs of AF647-labeled TOM20 in U2OS cells using 4Pi-BRAINSPOT (orange) and *in-vitro* method (black). Random aberrations were introduced via deformable mirrors for each trial. Box is the upper and lower quartiles, central line is the median, and error bar is the furthest data point excluding outliers. Statistical significance: t-test.

We further evaluated 4Pi-BRAINSPOT experimentally with induced aberrations where 4Pi-BRAINSPOT is used to retrieve coherent PSF models from the measured data where independent wavefront distortions are generated by deformable mirrors in both interferometric arms. Among all 7 types of commonly encountered aberrations, represented by Zernike polynomials (Figure S1A-H; Method 1.10), *in-situ* 4Pi-PSF models achieved high similarity compared to the recorded ones (NCC of 0.98 ± 0.01, n=14 trials), a significant improvement over their *in-vitro* or theoretical counterparts (NCC of 0.81 ± 0.04, n=84 trials). Our method also achieved an axial precision of 2-3 nm in a range of ±800 nm when localizing fluorescent beads, demonstrating the stability of the interferometric system (Figures S1I&J). Next, we assessed the capability of 4Pi-BRAINSPOT algorithm to capture dynamic 4Pi distortions while imaging immunofluorescence-labeled TOM20 in U2OS cells. To characterize the reliability of the time-dependent model update, we introduced random aberrations via deformable mirrors at various time points during imaging (Method 1.11). We evaluated the goodness of fitting between the captured emission patterns in cells and the in situ retrieved PSF models by calculating the log-likelihood ratio (LLR) (Figure 2I; Method 3.7). Our method demonstrated an LLR of 1384 ± 208 across 20 trials with various aberrations, consistently outperforming the conventional static *in-situ* model constructed from bead measurement near the cells with an LLR of 1769 ± 298 (n=200,218 localizations).

### Nanoscale visualization and quantitative analysis of subcellular organisms in mammalian cells

To demonstrate the capabilities of 4Pi-BRAINSPOT in resolving subcellular ultrastructure, we imaged microtubular structure immunofluorescence-labeled with α-tubulin in COS-7 cells, known to be 3D tubules with diameters of 50-70 nm ^38,46,47^, on top coverslips with a 25-μm cavity, as a controlled aberrated condition (Figure 3A; Method 1.6). The inhomogeneous refractive indices of the specimen, water-based mounting medium, and objective immersion oil cause refractive index mismatch aberrations and a random change in the cavity phase. These, in turn, exacerbate errors when pin-pointing the center localizations of single emitters through thick specimens due to the mismatch between the captured emission pattern and the *in-vitro* bead model. We compared the result of the *in-situ* model obtained from 4Pi-BRAINSPOT with that of two *in-vitro* modeling methods—voxel-based and pupil-based modeling ^36,37^— using bead measurements taken from both the top coverslip near the structures of interest and the bottom coverslip away from the structures (Figures 3N&O). With 4Pi-BRAINSPOT, we found that the expected hollow tubular structures of microtubules could be revealed as distinct ring cross-sections at various positions and depths (Figures 3B-M). In comparison, the result of 4Pi-BRAINSPOT illustrated sharp contours, whereas the other methods exhibited loose distributions or high localization variability (Figure 3P).

**Figure 3.**
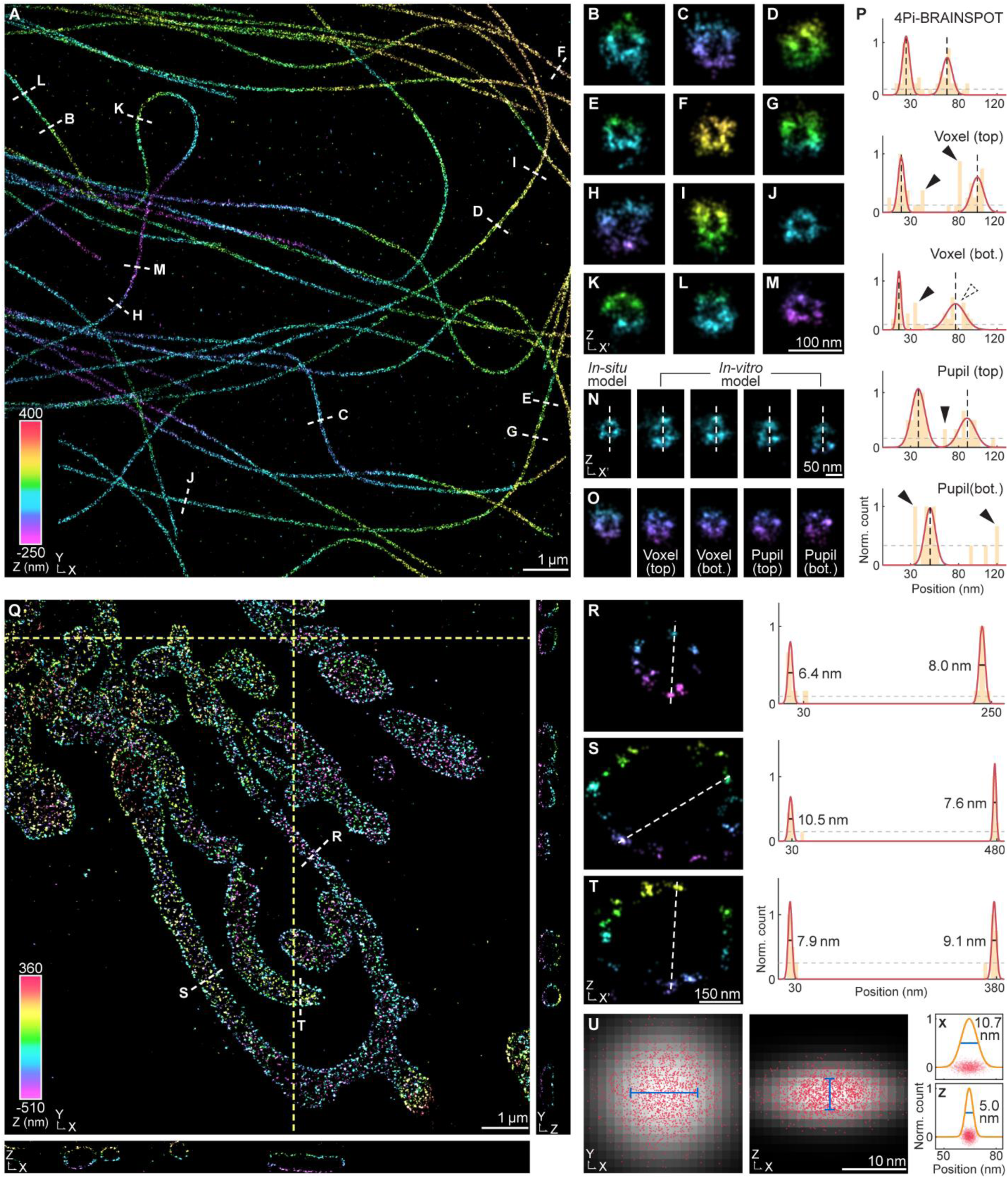
Performance of 4Pi-BRAINSPOT in mammalian cells. (**A**) Reconstruction of AF647-labeled α-tubulin in COS-7 cells with 25-µm cavity, color-coded according to single-molecule axial positions. (**B**-**M**) Vertical cross-sections of selected positions (white dashed lines) in (**A**). (**N**&**O**) Comparisons of vertical cross-sections between 4Pi-BRAINSPOT, *in-vitro* voxel-based method using beads on top coverslips (Voxel (top)), *in-vitro* voxel-based method using beads on bottom coverslips (Voxel (bot.)), *in-vitro* pupil-based method using beads on top coverslips (Pupil (top)), and *in-vitro* pupil-based method using beads on bottom coverslips (Pupil (bot.)). (**P**) Normalized intensity profile along white dashed lines in (N). Yellow bar is the localization histogram with 6-nm bins and red curve is the fitted Gaussian function to the histogram. Features isolated or below the exclusion threshold (gray dashed line) are excluded from analysis. Black dashed line marks the estimated feature center, solid arrowhead indicates excluded isolated features, and dashed arrowhead indicates feature discontinuity. (**Q**) Reconstruction of AF647-labeled TOM20 in COS-7 cells with 25-µm cavity with horizontal and vertical cross-sections of selected positions (yellow dashed lines). (**R**-**T**) Vertical cross-sections (left) of targeting mitochondria (white dashed lines) in (**Q**). Normalized intensity profile (right) along white dashed lines in (**R**-**T**). Yellow bar is the localization histogram with 6-nm bins, red curve is fitted Gaussian functions to the histogram, gray dashed line is the exclusion threshold, and black solid line marks the estimated feature FWHM. (**U**) Horizontal (left) and vertical (middle) cross-sections and normalized intensity profiles (right) of superposed single-molecule cluster. Red dot is single-molecule localization (n=1845 localizations), grayscale distribution is rendered cluster using 2D Gaussian blur with estimated FWHM, orange curve is fitted Gaussian function, and blue line marks estimated FWHM. See also Figure S4. See also Figure **S5**.

Next, we imaged mitochondria immunofluorescence-labeled with TOM20 in COS-7 cells, a receptor protein clustering on the outer mitochondrial membrane ^48–50^, through a 25µm water-based cavity (Figure 3Q). We successfully visualized the nanoscale distribution of the TOM20 clusters on the mitochondrial surface, achieving a full-width half maximum (FWHM) of 6-10 nm (Figures 3R-T). To validate the achieved molecular localization precision *in situ*, we measured the sizes of localization distribution formed by overlapping 3D-aligned single-molecule localization clusters (Figure S5; Method 4.5). These measurements aligned with the Cramér-Rao lower bound (CRLB) resolution of 11.3±3.8 nm laterally and 4.7±1.5 nm axially (corresponding to localization precision of 4.8±1.6 nm and 2.0±0.6 nm, n=1845 localizations across 387 clusters; Method 3.8).

Furthermore, we examined the organization of immunofluorescence-labeled Reticulon 4 (Rtn4), a protein stabilizing the curvature of the endoplasmic reticulum (ER) membrane, in COS-7 cells (Figure 4A) ^51,52^. Despite various hypotheses about Rtn4’s role were suggested, traditional microscopy has been limited in providing nanoscale 3D insights into Rtn4 regulation in the ring closure process ^53–56^. Using 4Pi-BRAINSPOT, we observed two distinct patterns of Rtn4-enriched ER membranes (Figure 4B) ^52,57^: narrow-waist ring structures about 30-nm radius (Figures 4C&D, H), indicative of ER tubules with small round lumens, and parallel lines separated by 100 nm (Figures 4E-G, I), suggesting ER tubules with larger elliptical lumens. The result consisting of high-precision molecular localizations in 3D allowed us to numerically flatten the ER membrane, map the angular distribution of Rtn4 clusters, and visualize the evolving spatial arrangement and transition of Rtn4 along an isolated ER segment (Fig. 4K-M).

**Figure 4.**
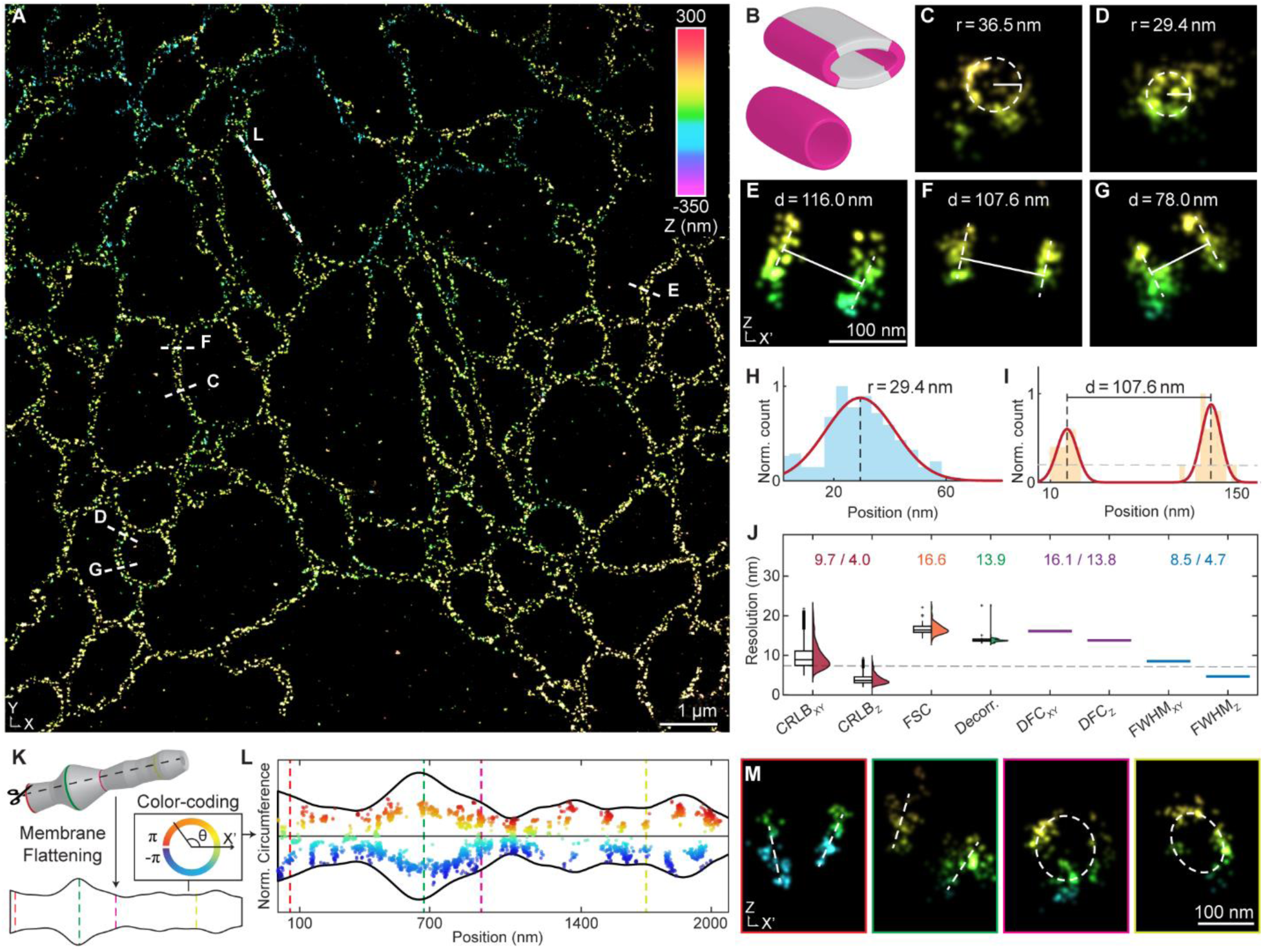
Demonstration of endoplasmic reticulum (ER) tubules in mammalian cells using 4Pi-BRAINSPOT (**A**) Reconstruction of AF647-labeled Rtn-4 in COS-7 cells, color-coded according to single-molecule axial positions. (**B**) Illustration of ER tubules with elliptical lumens (top), where Rtn-4 (magenta) concentrates on the high-curvature edges only, and round lumens (bottom), where Rtn-4 distributed across the surface. (**C**&**D**) Vertical cross-sections of selected positions (white dashed lines) in (**A**) showing ring structures. White dashed circle marks the estimated ring structure and white solid line indicates its radius. (**E**-**G**) Vertical cross-sections of selected positions (white dashed lines) in (**A**) showing parallel lines. White dashed circle marks estimated double-line structure and white solid line indicates interval distance. (**H**) Radius analysis in (**D**). Blue bar is the distance histogram between localizations and the structural center with 4-nm bins, red curve is fitted Gaussian functions to the histogram, and black dashed line marks the estimated radius. (I) Interval distance analysis in (**F**). Yellow bar is the localization histogram with 6-nm bins, red curve is fitted Gaussian functions to the histogram, gray dashed line is the exclusion threshold, black dashed line marks the estimated feature center, and black solid line indicates the estimated interval distance. (**J**) Comparison of resolution analysis using different methods, including lateral and axial CRLB-estimated resolutions (CRLB_XY_ of 9.7±3.1 nm and CRLB_Z_ of 4.0±1.3 nm, n=2955 localizations), FSC resolution (FSC of 16.6±1.3 nm, n=81 subregions), imaging decorrelation resolution (Decor. of 13.9±1.0 nm, n=81 subregions), lateral and axial DFC resolutions (DFC_XY_ of 16.1 nm and DFC_Z_ of 13.8 nm), and measured lateral and axial FWHMs of superposed single-molecule cluster (FWHM_XY_ of 8.5 nm and FWHM_Z_ of 4.7 nm). Box is the upper and lower quartiles, central line is the median, error bar is the furthest data point excluding outliers, gray dashed line is the estimated isotropic 3D CRLB-estimated resolution, and the distribution is shown on the right side. (**K**) Workflow of circumference analysis. We first sliced tubular structures along the longitudinal direction and flattened them as 2D surfaces. Localizations are then color-coded according to angular positions ranging from ±π. (**L**) Circumference analysis of targeting ER tubule in (A). (**M**) Vertical cross-sections of selected positions (corresponding colors) in (**L**), showing varied distributions of Rtn-4 from double-line (red and green) to ring (magenta and yellow) structures.

### 4Pi-BRAINSPOT resolves ultrastructure of dendrites and spines through brain tissues

Dendritic spines are crucial for synaptic connections and play a key role in memory processing by reflecting the brain’s adaptive capacity through their morphology changes driven by synaptic plasticity. Synaptic potentiation can prompt the formation of new spines or alter the concavity of the spine head and the dimensions of the spine neck. Thin spines are often linked to learning and memory, mushroom-shaped spines to mature synaptic connections, and stubby spines to developmental or repair processes ^58–60^. Furthermore, irregularities in spine organization and density are also associated with synaptic connectivity disruptions ^61–63^. Understanding these nanoscale changes in spines is crucial for elucidating the mechanisms of synaptic connection and pathology of neurological disorders.

Channelrhodopsin-2 (ChR2), a light-gated ion channel, serves as an important role for investigating optogenetic modulation ^64–67^. Its distribution and clustering within the neuronal membrane makes it an ideal target for assessing the imaging fidelity of our method. To visualize the spatial distribution of these ion channels, we imaged 50-μm mouse brain slices and successfully located the distinct membrane-associated protein clusters throughout dendritic spines. Our quantitative resolution analysis indicated an achieved sub-15 nm resolution, and the resulting reconstruction revealed the 3D molecular distribution of ChR2 on dendritic spines (Figures 5A&6A). We were able to observe structural transitions from dendritic shafts to spine heads, where the 3D localized ChR2 molecular cloud provided detailed morphological information. We also observed a distinct concave spine head on a long and mushroom-shaped spine, potentially indicating an increased postsynaptic density area in response to synaptic adaptation ^68^.

**Figure 5.**
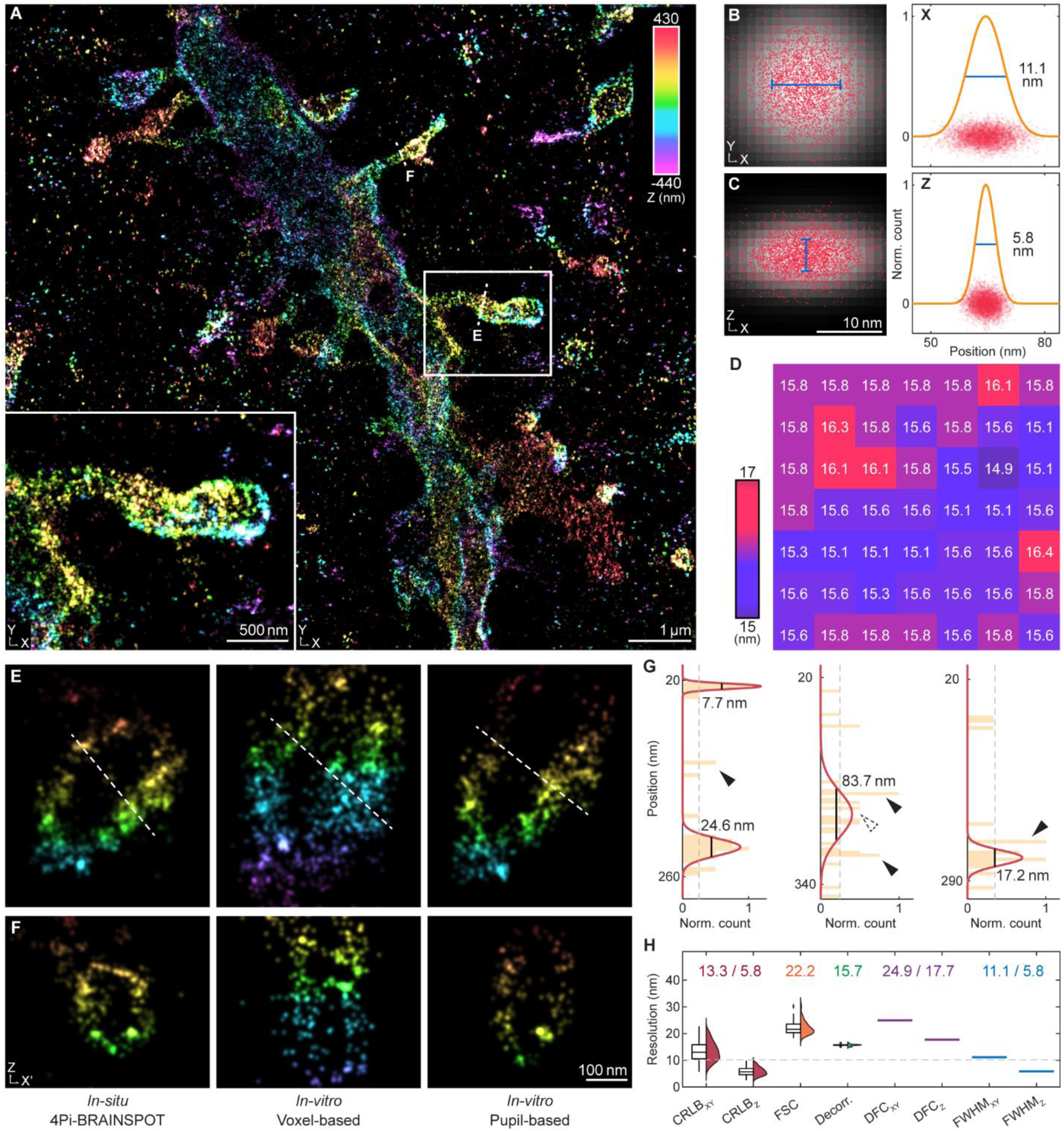
Demonstration of 4Pi-BRAINSPOT in transgenic mouse brain slices. (**A**) Reconstruction of AF647-labeled ChR2 in 50-µm thick transgenic mouse brain slices using 4Pi-BRAINSPOT, color-coded according to single-molecule axial positions. Inset shows enlarged dendritic spines in white solid box. (**B**&**C**) Horizontal (**B**) and vertical (**C**) cross-sections (left) and normalized intensity profiles (right) of superposed single-molecule cluster. Red dot is single-molecule localization (n=4272 localizations), grayscale distribution is rendered cluster using 2D Gaussian blur with estimated FWHM, orange curve is fitted Gaussian function, and blue line marks estimated FWHM. (**D**) Heatmap of image decorrelation resolutions by segmenting 2D projection of (**A**) into 7×7 subregions. (**E**&**F**) Vertical cross-sections of targeting dendritic spines (white dashed lines) in (**A**) using 4Pi-BRAINSPOT (left), *in-vitro* voxel-based method (middle), and *in-vitro* pupil-based method (right). (**G**) Normalized intensity profile along white dashed lines in (**E**). Yellow bar is the localization histogram with 6-nm bins, red curve is fitted Gaussian functions to the histogram, gray dashed line is the exclusion threshold, black solid line marks the estimated feature FWHM, solid arrowhead indicates excluded isolated features, and dashed arrowhead indicates feature discontinuity. (**H**) Comparison of resolution analysis using different methods, including lateral and axial CRLB-estimated resolutions (CRLB_XY_ of 13.3±2.6 nm and CRLB_Z_ of 5.8±1.5 nm, n=4272 localizations), FSC resolution (FSC of 22.2±3.6 nm, n=81 subregions), imaging decorrelation resolution (Decor. of 15.7±0.4 nm, n=81 subregions), lateral and axial DFC resolutions (DFC_XY_ of 24.9 nm and DFC_Z_ of 17.7 nm), and measured lateral and axial FWHMs of superposed single-molecule cluster (FWHM_XY_ of 11.1 nm and FWHM_Z_ of 5.8 nm). Box is the upper and lower quartiles, central line is the median, error bar is the furthest data point excluding outliers, gray dashed line is the estimated isotropic 3D CRLB-estimated resolution, and the distribution is shown on the right side. See also Figure **S7**.

**Figure 6.**
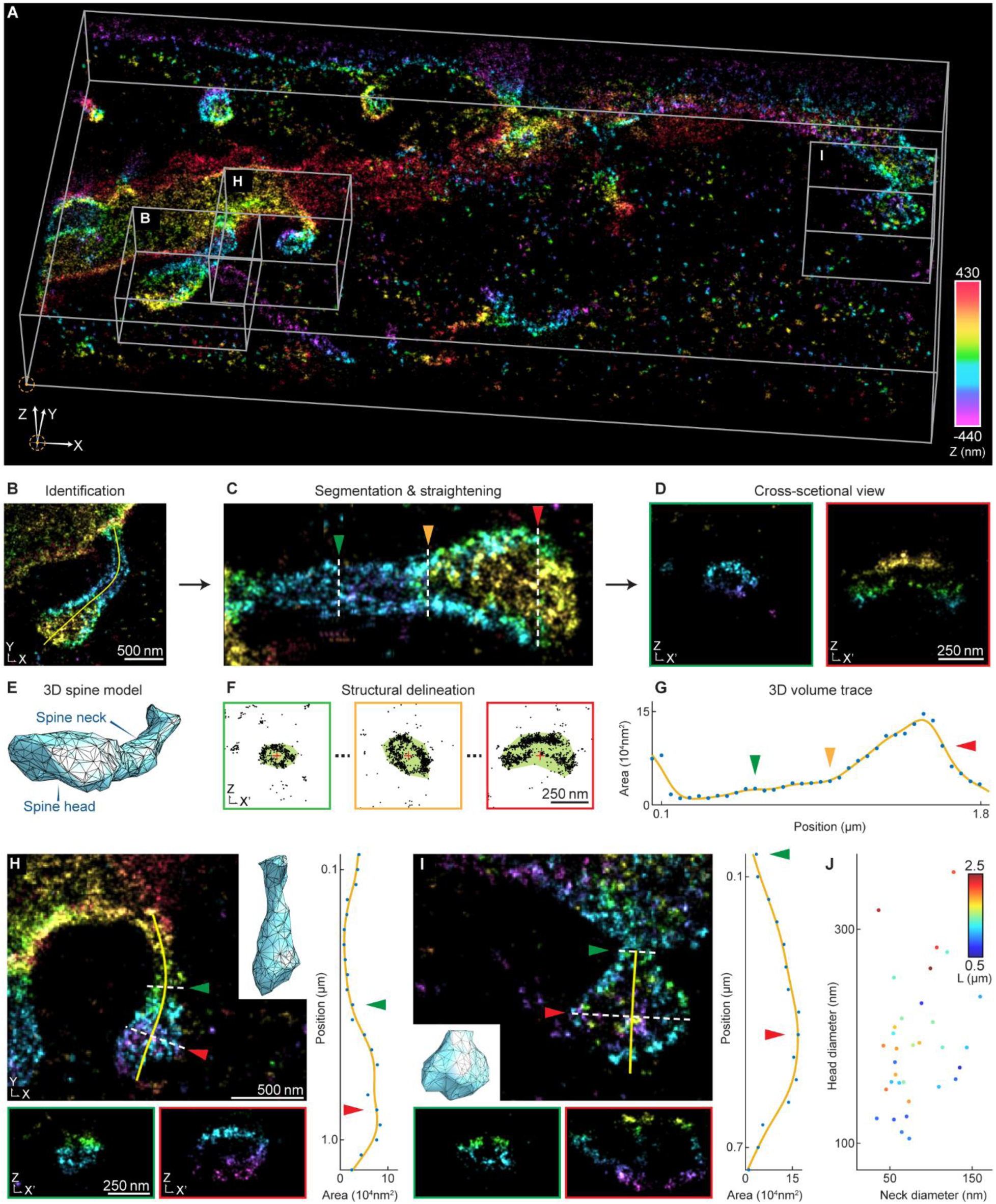
Demonstration of spine morphological analysis. (**A**) Reconstruction of AF647-labeled ChR2 in 50-µm thick transgenic mouse brain slices using 4Pi-BRAINSPOT, color-coded according to single-molecule axial positions. (**B**-**G**) Workflow of analysis pipeline of 3D structural morphology. (**B**) Identification of targeting spine (white boxes) in (**A**). Yellow curve is the trace of algorithm from manual identification. (**C**) Straighten structure of the dendritic spine in (**B**) by segmenting localizations into 150-nm segments with 50-nm overlapping. (**D**) Vertical cross-sections of selected positions (white dashed lines with colored arrowheads) in (**C**) showing spine neck (green) and spine head (red). (**E**) 3D model of the spine in (**B**). (**F**) Localization distribution at selected positions (white dashed lines with colored arrowheads) in (**C**). Black dot is localization, green area marks the alpha-shaped boundary, and red cross marks the estimated structural center. (G) Volume quantification of dendritic spine in (**C**). Blue dot is the measured area and yellow curve is the fitted curve. Colored arrowhead indicates the area corresponding in (**C**). (**H**&**I**) Demonstrations of spine analysis of targeting spines (white boxes) in (**A**) with volume quantification (right). Inset is a 3D model of the spine, green and red box are vertical cross-sections of selected positions of the spine neck and head pointed by colored arrowheads. (**I**) Distribution of diameters of spine necks and spine heads (n=33 spines, N=16 tissues), color-coded according to spine length.

To assess the performance of 4Pi-BRAINSPOT through brain tissues, we evaluated the resolution with multiple criteria. Our cluster analysis pipeline indicated that the overall size of single-molecule clusters distributed through different tissues was 13.0±2.2 nm laterally and 6.8±1.3 axially (n=7 datasets; Figures 5B, 5C, S7F, and S7G; Method 4.5). We verified it by measuring the FWHM of the neuronal membrane waist as ∼8 nm (Figures 5G, S7B, and S7C). These measurements are consistent with the CRLB-estimated resolution of 15.1±2.6 nm laterally and 7.0±1.6 axially. To evaluate the achievable resolution under certain labeling density and imaging conditions, we utilized directional Fourier correlation (DFC) analysis to calculate Fourier correlation resolution in spatial-frequency domains. The DFC results indicated lateral and axial resolutions of 30.3±5.8 nm and 21.0±4.7 nm, closely matching the results from 2D imaging decorrelation analysis of 17.8±3.5 nm (Figures 5D&S7D) and 3D Fourier shell correlation of 26.6±5.0 nm (n=7 datasets; Figure 5H&S7E; Method 4.6).

Furthermore, we developed a tailored pipeline for processing 4Pi-SMSN data to quantitatively analyze dendritic spine morphology based on the resulting molecular coordinates of individual membrane-bounded proteins in 3D (Figure 6B-G; Method 4.1). This method, first, segments single molecule localizations into sequential cross-sections, then detects the complex boundaries of major structures from scattered molecular centers using the alpha-shape algorithm, which unites multiple convex hulls encompassing localization subsets and finally pinpoints structural centers by fitting ellipses to localizations within the alpha-shaped boundaries. The radii of the isolated spine sections are then estimated by calculating the mode of distances between the center and the localizations. Using the automated quantification pipeline, we analyzed the ChR2-outlined membrane contours and quantified the topographic and volumetric changes of spines (Figures 6H&I). In a collection of 33 identified spines imaged at the visual cortex resolved from brain tissues (Figure 6J), we observed a diverse range of spine morphologies, from 0.6-µm-long round-shaped spines to 2.4-µm-long mushroom-shaped spines. The spine neck radii varied from 34.1 to 159.7 nm (81.1 ± 33.6 nm), and spine heads radii varied from 104.3 to 353.3 nm (193.1 ± 59.3 nm), with total volumes ranging from 0.02 to 0.36 µm^3^ (0.10 ± 0.07 µm^3^), highlighting the effectiveness of our method in analyzing spines with diverse arrangements and orientations.

## Discussion

Understanding the brain has long been one of the most ambitious and compelling endeavors in science. However, a grand challenge remains in neuroscience—visualizing how sensory experiences sculpt neural circuits and synaptic connections at the subcellular level has been limited due to the challenges of achieving sufficient resolution and molecular specificity deep within the brain. While previous research has made significant contributions to understanding the molecular and cellular mechanisms underlying neural plasticity, current super-resolution imaging techniques have mostly been limited to cultured preparations. Despite 2-photon imaging being widely used in live animals, its intrinsic diffraction limit prevents it from capturing fine structures and protein localizations deep within neural tissues ^69,70^. 4Pi-BRAINSPOT bridges this profound gap by stepping forward in achievable resolution and depth, enabling deciphering nanoscale molecular architecture deep within the brain tissues with unparalleled clarity. Furthermore, the highly accurate 3D molecular map generated by 4Pi-BRAINSPOT supports a novel analysis pipeline that identifies, segments, and quantifies the morphological properties of dendritic spines. This capability facilitates advanced studies of synaptic architecture and plasticity, fundamental processes underlying learning, memory, and information processing.

Further enhancement for the 4Pi-BRAINSPOT system involves increasing throughput volume and addressing potential field-dependent distortions. When imaging a large FOV, these lateral-dependent distortions may alter the 4Pi-PSF, and thus challenge the assumption of shift-invariant PSF ^71,72^. To mitigate these issues, one might integrate field lenses specifically designed to compensate for FOV-dependent distortions at the instrumental level and develop algorithms to adapt to these variations. Implementing these improvements will ensure that *in-situ* modeling accuracy is maintained across the extended FOVs, and thus its localization performance for high-fidelity, high-throughput imaging applications. Integrating functional imaging techniques will also enhance 4Pi-BRAINSPOT’s ability to elucidate the molecular mechanisms underlying neural activities ^29,31^. Utilizing 4Pi-BRAINSPOT in thick brain tissue to study the perceptual experience-dependent plasticity of dendritic spines can correlate with the functional synaptic measurements by electrophysiology experiment. This would enable nanoscale molecular exploration, providing a comprehensive understanding of how structural changes at the nanoscale level impact overall neuronal function and behavior.

We believe 4Pi-BRAINSPOT marks a substantial advancement in advanced optical nanoscopy, offering unprecedented insights into the nanoscale organization of biological tissues. We hope this unique observation capacity near the molecular level will further our understanding of cellular functions, neural circuits, and disease pathologies.

## Acknowledgment

We thank Peiyi Zheng (Purdue University) for suggestions on the super-resolution reconstruction process and microscope optimization. We thank Sheng Liu and Vamara Dembele (Purdue University) for their suggestions on microscope optimization. We thank George Takahashi and Vaughn M Valentino (Envision Center at Purdue University) for helping with the 3D visualization of the dataset. This work was supported by the US National Institutes of Health (grants GM119785 to F.H., MH123401 to F.H. and A.A.C.). F.X. acknowledges the National Key R&D Program of China (2023YFC3402600) and the National Natural Science Foundation of China (62272041).

## Author contributions

Conceptualization, H.C.G., F.X., and F.H.; Methodology, H.C.G. and F.X.; Software, F.X., Y.L., T.C., and Y.L.; Validation, H.C.G. and F.X.; Formal Analysis, H.C.G. and F.X.; Investigation, H.C.G., X.C., and A.A.C.; Resources, X.C., C.B., Y.Z., A.A.C., and F.H.; Visualization, H.C.G. and F.X.; Funding Acquisition, F.X. A.A.C., and F.H.; Supervision, F.X. A.A.C., and F.H.; Writing, all authors.

