## Supplementary Information for "Interferometric Ultra-High Resolution 3D Imaging through Brain Sections"

This PDF file includes:

Supplementary Figures S1 to S8

Supplementary Methods

Additional References

**Supplementary figures**

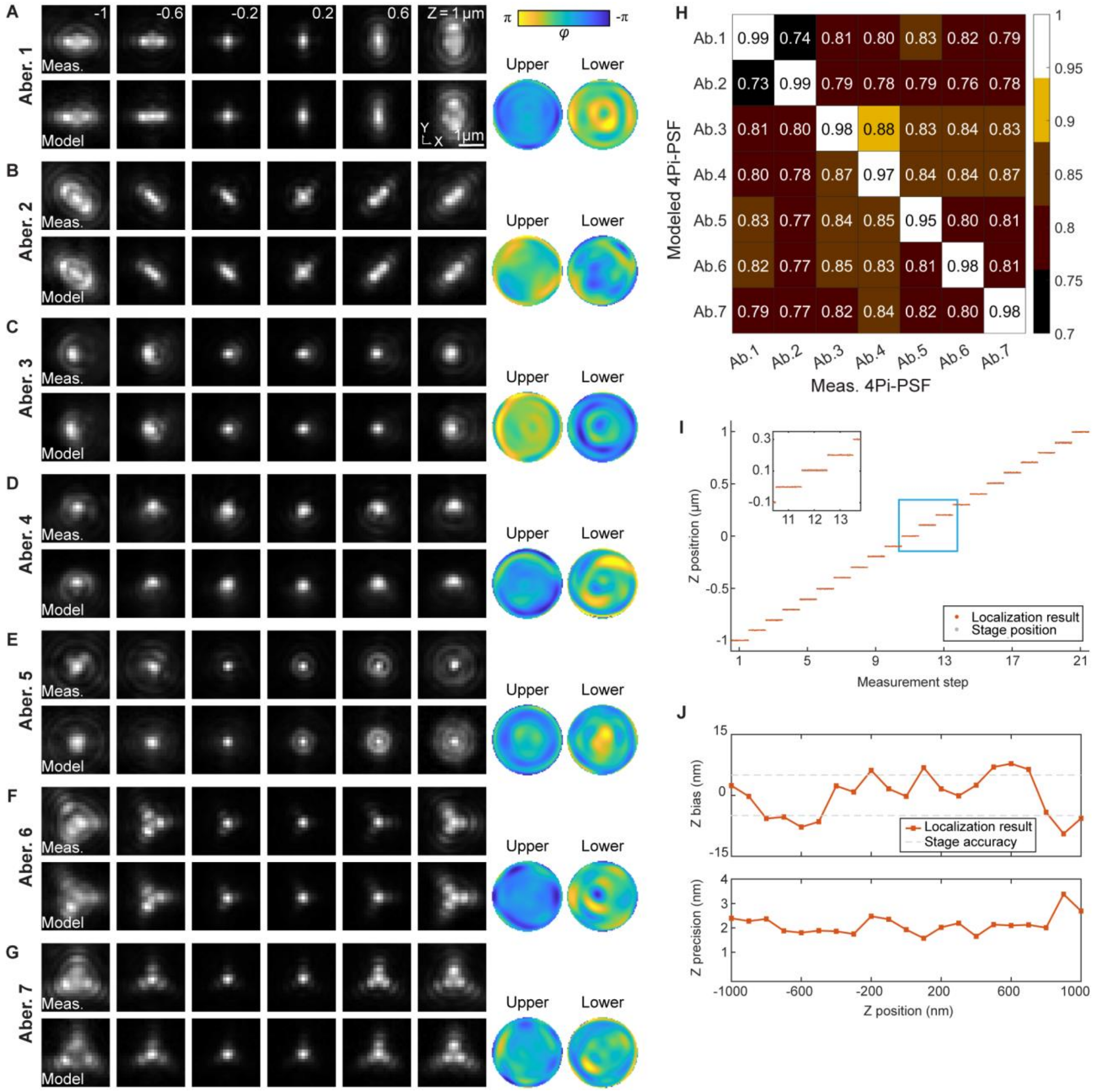


**Figure S1 Performance of *in-situ* PSF retrieval. Related to Figure 1.**

(**A**-**G**) Interferometric pattern comparison between fluorescent bead measurements (top row) and 4Pi-BRAINSPOT PSF models (bottom row) in channel p1 with different aberrations at axial positions ranging from ±1 μm. Retrieved coherent pupil functions are shown on the right.

(**H**) Heatmap of NCC similarity between measured and modeled 4Pi-PSFs in (**A**-**G**).

(**I**) Stepwise localizations of bead measurements with 100 nm increments and 50 measurements per step over an axial range from ±1 μm using 4Pi-BRAINSPOT. Orange dot is localization result of 4Pi-BRAINSPOT, and gray dot is assigned stage position. Inset shows enlarged result in cyan box.

(**J**) Estimation of bias (top row) by calculating mean distances of localization results from assigned stage positions and precision (bottom row) by calculating standard deviations of localization results in (**I**). Orange curve is localization result, and gray dashed line is the stage accuracy.

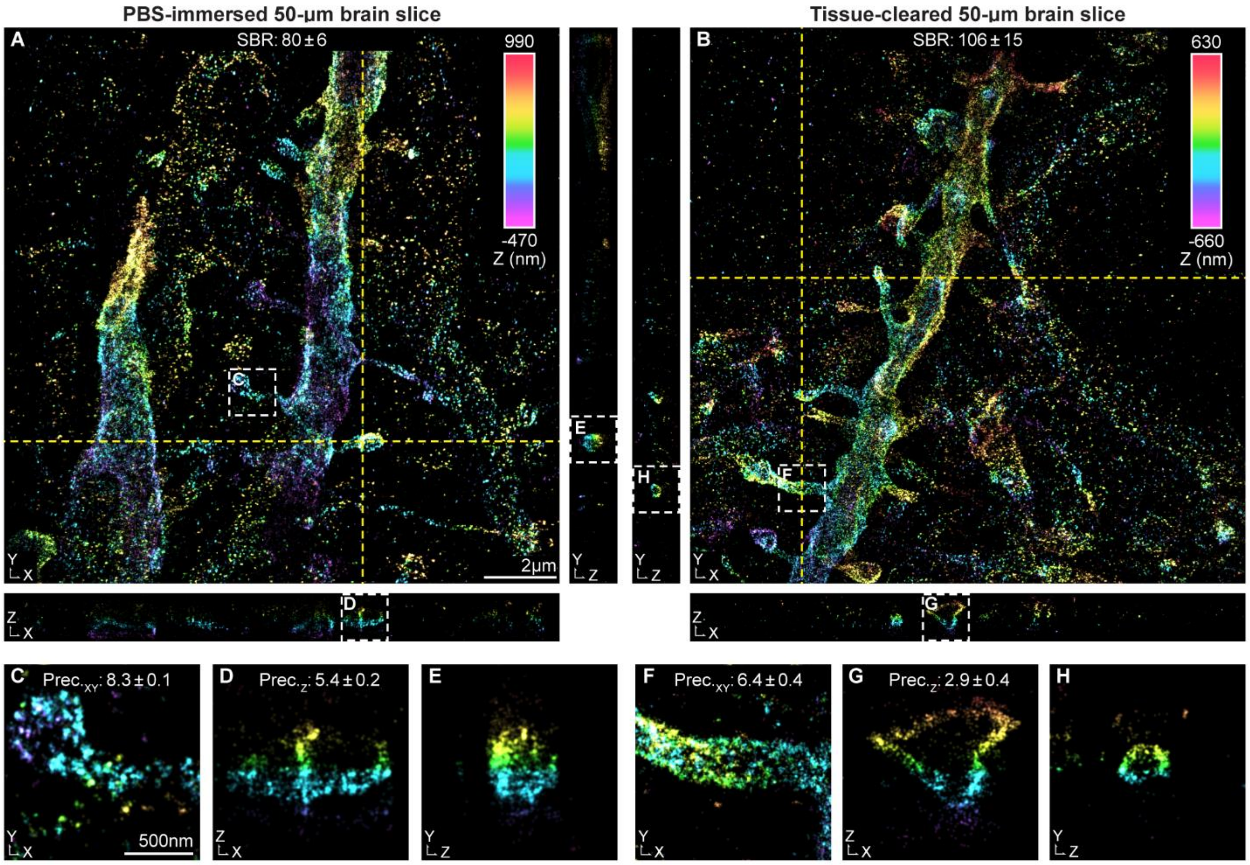


**Figure S2 Comparative analysis of 4Pi-BRAINSPOT in mouse brain slices with and without tissue clearing process. Related to Figure 1.**

(**A**&**B**) Reconstruction and cross-sections of selected positions (yellow dashed line) of AF647-labeled ChR2 in PBS-immersed (**A**) and tissue-cleared (**B**) 50-µm thick mouse brain slices using 4Pi-BRAINSPOT, color-coded according to the single-molecule Z positions. Signal-to-background ratio (SBR) of tissue-cleared specimens was 106±15 (n=12 datasets) and that of untreated specimens was 80±6 (n=6 datasets), respectively.

(**C**-**H**) Enlarged views of selected regions (white dashed boxes) in (**A**&**B**). The lateral and axial localization of tissue-cleared specimens are 6.4±0.4 and 2.9±0.4 (n=12 datasets), while those of untreated specimens are 8.3±0.1 and 5.4±0.2 (n=6 datasets).

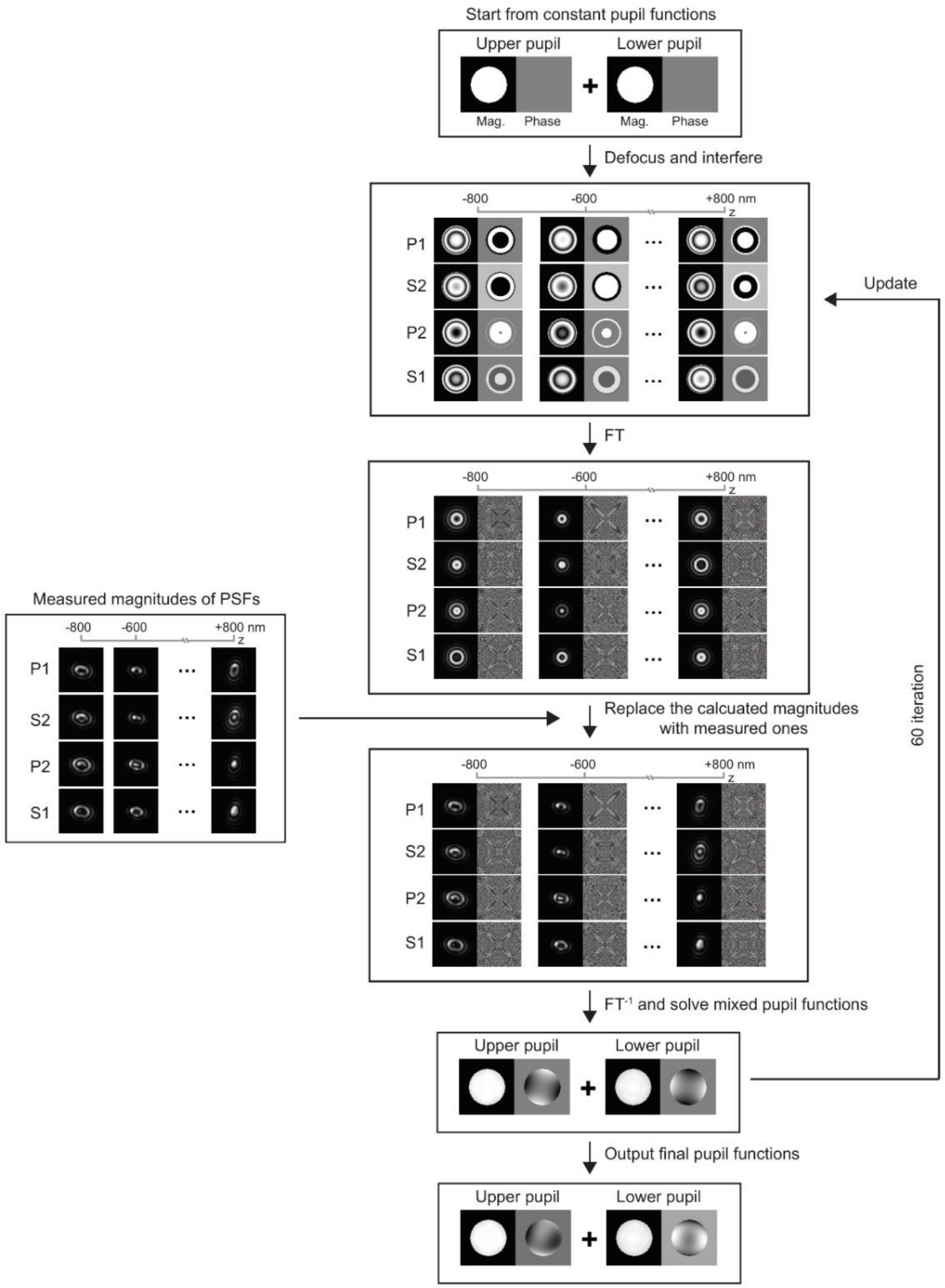


**Figure S3 Workflow of coherent 4Pi phase retrieval process. Related to Figure 1.**

Coherent 4Pi phase retrieval began with initial coherent pupil functions with predetermined parameters. Superposed wavefields were generated by adding variable defocuses and phase delays across channels at axial positions ranging from ±800 nm. Amplitudes of 4Pi-PSF were computed by Fourier transform of superposed wavefield. Next, calculated magnitudes were replaced with the square root of obtained values, while maintaining original phases. Inverse Fourier transform was used to update superposed wavefields after which coherent pupil functions were updated by Moore–Penrose pseudo-inverse process. Results after 60 iterations were regarded as coherent pupil functions of obtained interferometric patterns.

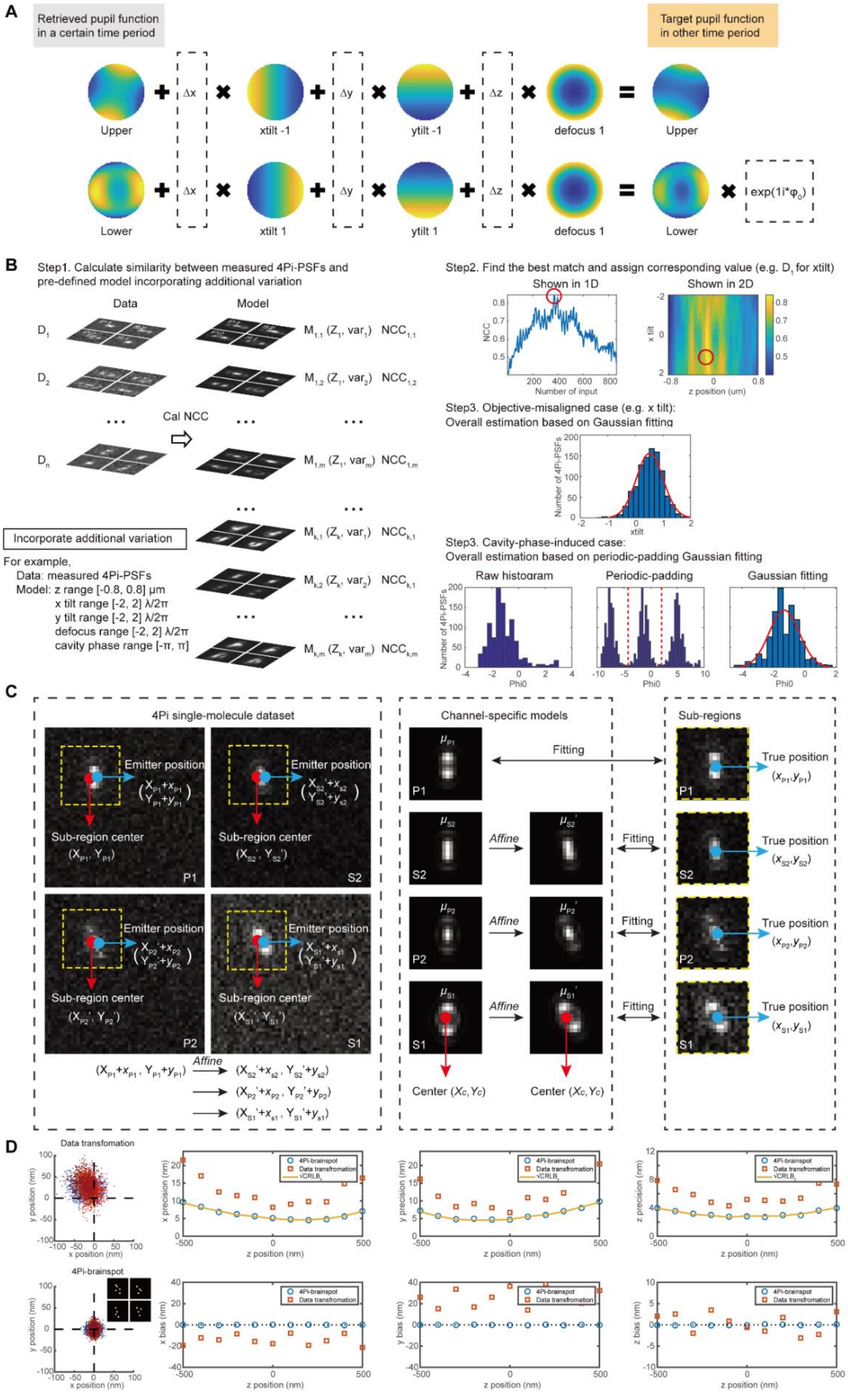
A
Retrieved pupil function in a certain time period
Target pupil function in other time period
Δx
Δy
Δz
Upper
xtilt -1
ytilt -1
defocus 1
Upper
Lower
xtilt 1
ytilt 1
defocus 1
Lower
exp(1i*φ0)
B
Step1. Calculate similarity between measured 4Pi-PSFs and pre-defined model incorporating additional variation
Data
Model
D1
D2
Dn
Cal NCC
M1,1 (Z1, var1) NCC1,1
M1,2 (Z1, var2) NCC1,2
M1,m (Z1, varm) NCC1,m
Mk,1 (Zk, var1) NCCk,1
Mk,2 (Zk, var2) NCCk,1
Mk,m (Zk, varm) NCCk,m
Incorporate additional variation
For example,
Data: measured 4Pi-PSFs
Model: z range [-0.8, 0.8] μm
x tilt range [-2, 2] λ/2π
y tilt range [-2, 2] λ/2π
defocus range [-2, 2] λ/2π
cavity phase range [-π, π]
Step2. Find the best match and assign corresponding value (e.g. D1 for xtilt)
Shown in 1D
Shown in 2D
NCC
Number of input
x tilt
z position (um)
Step3. Objective-misaligned case (e.g. x tilt):
Overall estimation based on Gaussian fitting
Number of 4Pi-PSFs
xtilt
Step3. Cavity-phase-induced case:
Overall estimation based on periodic-padding Gaussian fitting
Raw histogram
Periodic-padding
Gaussian fitting
Number of 4Pi-PSFs
Phi0
C
4Pi single-molecule dataset
Emitter position
Sub-region center
(XP1, YP1)
(XS2', YS2')
(XP2', YP2')
(XS1', YS1')
P1
S2
P2
S1
(XP1+xP1, YP1+yP1)
Affine
(XS2'+xS2, YS2'+yS2)
(XP2'+xP2, YP2'+yP2)
(XS1'+xS1, YS1'+yS1)
Channel-specific models
Fitting
Affine
Center (Xc, Yc)
Sub-regions
True position
(xP1, yP1)
(xS2, yS2)
(xP2, yP2)
(xS1, yS1)
D
Data transfomation
4Pi-brainspot
x position (nm)
y position (nm)
4Pi-brainspot
Data transformation
√CRLB
x precision (nm)
y precision (nm)
z precision (nm)
x bias (nm)
y bias (nm)
z bias (nm)
z position (nm)

**Figure S4 Dynamic model update and channel-specific localization. Related to Figure 1.**

(**A**) Illustration of dynamic model update in each time window. 4Pi-BRAINSPOT estimated and accommodated time-dependent interferometric aberrations according to objective misalignments (ΔX, ΔY, and ΔZ) and cavity phase shift ($\varphi_0$).

(**B**) Data processing workflow of dynamic in-situ model update in 4Pi-BRAINSPOT. The algorithm segmented individual 4Pi emission patterns from single-molecule datasets to create a library. We generated a series of reference PSFs for each time window based on coherent pupil functions from the previous window, with adjustments of 3D objective misalignments (ΔX, ΔY, and ΔZ) and cavity phase shifts (φ0). For each reference, NCC similarity score between obtained patterns and reference 4Pi-PSFs are evaluated to determine the time-dependent parameters. For objective misalignments, the corresponding coefficients are determined by fitting the similarity scores with a Gaussian function, identifying the peak position. For cavity phase variation, the corresponding coefficients are determined by periodic-padding Gaussian fitting the similarity scores and identifying the peak position. Refined results are regarded as retrieved coherent pupil functions for a given time window.

(**C**) Data processing workflow of 3D channel-specific single-molecule localization. The process began with mapping coordinates in reference channel p1 into the other channels using calculated affine matrices. Channel-specific 4Pi-PSF models representing respective statistical properties of SCOMS are generated based on dynamic *in-situ* 4Pi-PSF model and used in further 4Pi localization.

(**D**) Comparison of localization bias and precision between 4Pi-BRAINSPOT using channel-specific 4Pi-PSF models (orange) and translating acquired patterns (blue) instead. Insets are the simulated variance distribution.

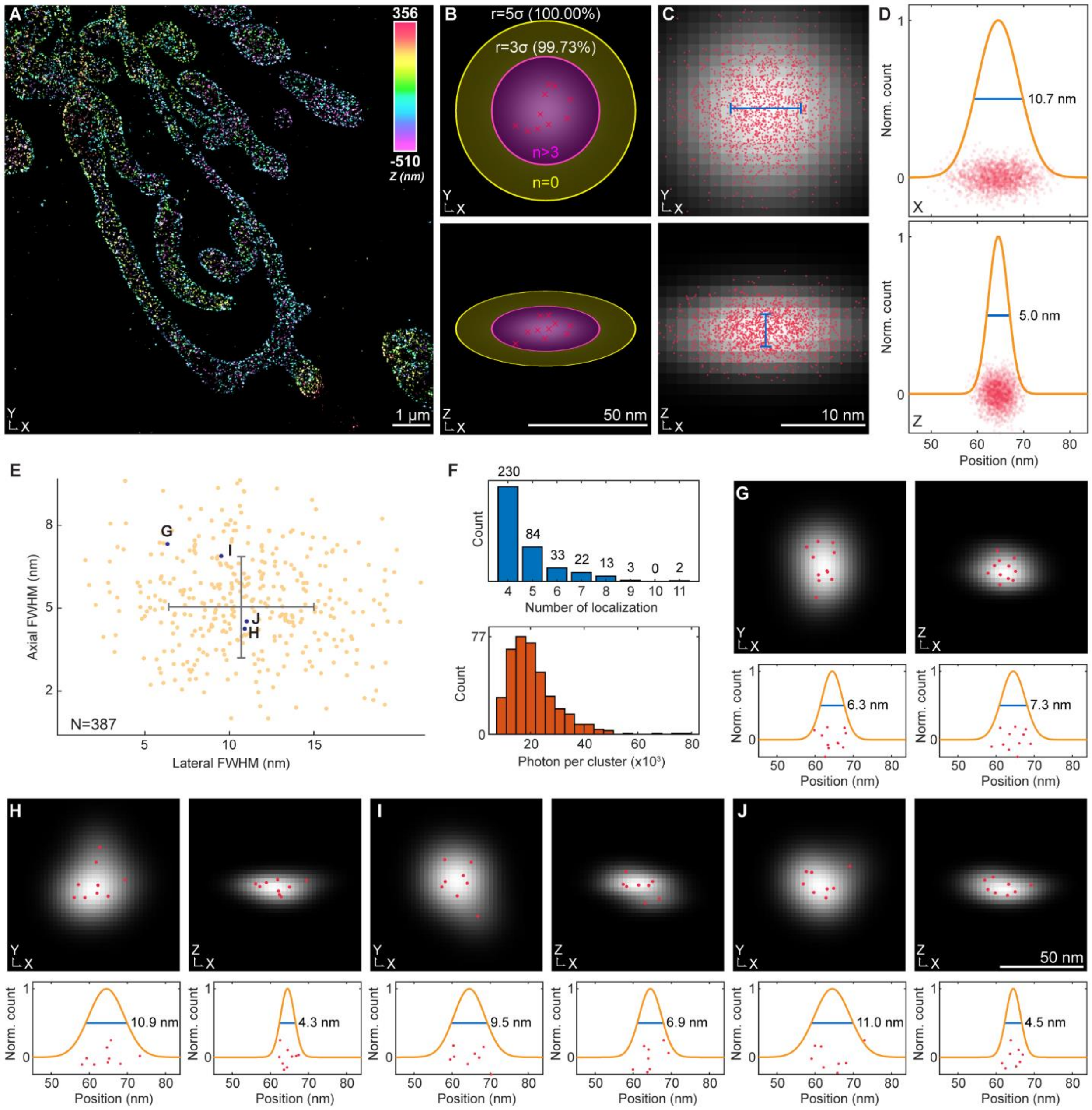


**Figure S5 Quantitative analysis of single-molecule localization clusters. Related to Figures 3 and 5.**

(**A**) Reconstruction of AF647-labeled TOM20 in COS-7 cells in Figure **3Q**, color-coded according to the single-molecule Z positions.

(**B**) Illustration of single-molecule cluster identification criteria. Magenta ellipse is region within three times of localization precision, yellow ellipse is region between three and five times of localization precision. Clusters were considered to originate from single molecules if more than three localizations fall in magenta region and none fall in yellow region. Red dot is simulated 7 localizations originated from single molecule and grayscale distribution is rendered cluster using 2D Gaussian blur.

(**C**) Cross-sectional views cluster superposition (n=3244 localizations). Red dots are single-molecule localizations, grayscale distributions are rendered images of localization cluster, and blue lines are FWHMs of localization distributions.

Horizontal (top) and vertical (bottom) cross-sections of superposed single-molecule cluster. Red dot is single-molecule localization (n=3244 localizations) and grayscale distribution is rendered cluster using 2D Gaussian blur with estimated FWHM.

(**D**) Horizontal (top) and vertical (bottom) normalized intensity profiles of superposed single-molecule cluster. Red dot is single-molecule localization (n=3244 localizations), orange curve is fitted Gaussian function, and blue line marks estimated FWHM.

(**E**) Lateral and axial FWHMs of all single-molecule clusters (10.3±3.0 and 5.3±1.9 nm, n = 387 clusters). Gray error bar indicates mean and standard deviation of lateral and axial FWHMs.

(**F**) Histograms of localization counts per cluster (top) and photon counts per cluster (bottom).

(**G**-**J**) Cross-sections and normalized intensity profiles for four selected clusters in (**E**) with >7 localizations each. Red dot is single-molecule localization (n=1845 localizations), grayscale distribution is rendered cluster using 2D Gaussian blur with estimated FWHM, orange curve is fitted Gaussian function, and blue line marks estimated FWHM.

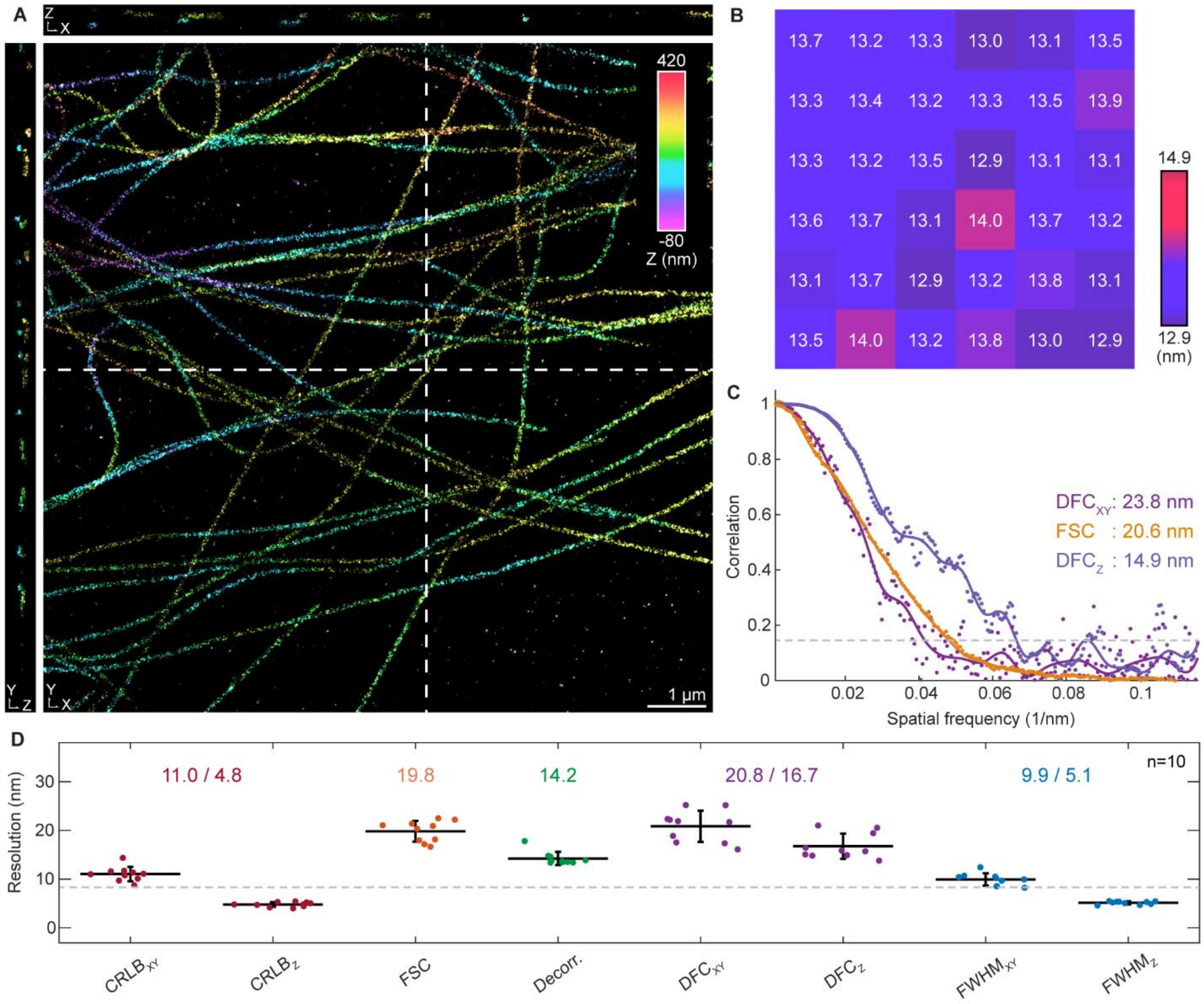


**Figure S6 Resolution assessment of 4Pi-BRAINSPOT in mammalian cells. Related to Figure 3.**

(**A**) Reconstruction and cross-sections of selected positions (white dashed lines) of AF647-labeled α-tubulin in COS-7 cells with a 25-μm cavity using 4Pi-BRAINSPOT, color-coded according to the single-molecule Z positions.

(**B**) Heatmap of image decorrelation resolutions by segmenting 2D projection of (**A**) into 6x6 subregions.

(**C**) Correlation analysis of Fourier shell correlation (FSC, orange) and directional Fourier correlation laterally and axially ($DFC_{XY}$, purple; $DFC_{Z}$, navy) at different spatial frequencies. Marker is correlation coefficient at selected spatial frequency, solid line is cubic-spine-fitted curve, and gray dashed line is 1/7 criteria to determine resolutions.

(**D**) Comparison of resolution analysis using different methods (n=10 datasets, including 4 datasets of AF647-labeled α-tubulin, 3 datasets of TOM20, and 3 datasets of Rtn4 in COS-7 cells), including lateral and axial CRLB-estimated resolutions ($CRLB_{XY}$ of 11.0±1.4 nm and $CRLB_{Z}$ of 4.8±0.4 nm, red), FSC resolution (FSC of 19.8±2.0 nm, orange), imaging decorrelation resolution (Decor. of 14.2±1.3 nm, green), lateral and axial DFC resolutions ($DFC_{XY}$ of 20.8±3.1 nm and $DFC_{Z}$ of 16.7±2.5 nm, purple), and measured lateral and axial FWHMs of superposed single-molecule cluster ($FWHM_{XY}$ of 9.9±1.2 nm and $FWHM_{Z}$ of 5.1±0.3 nm, blue). Central line is mean, error bar is standard deviations, and gray dashed line is estimated isotropic 3D CRLB-estimated resolution.

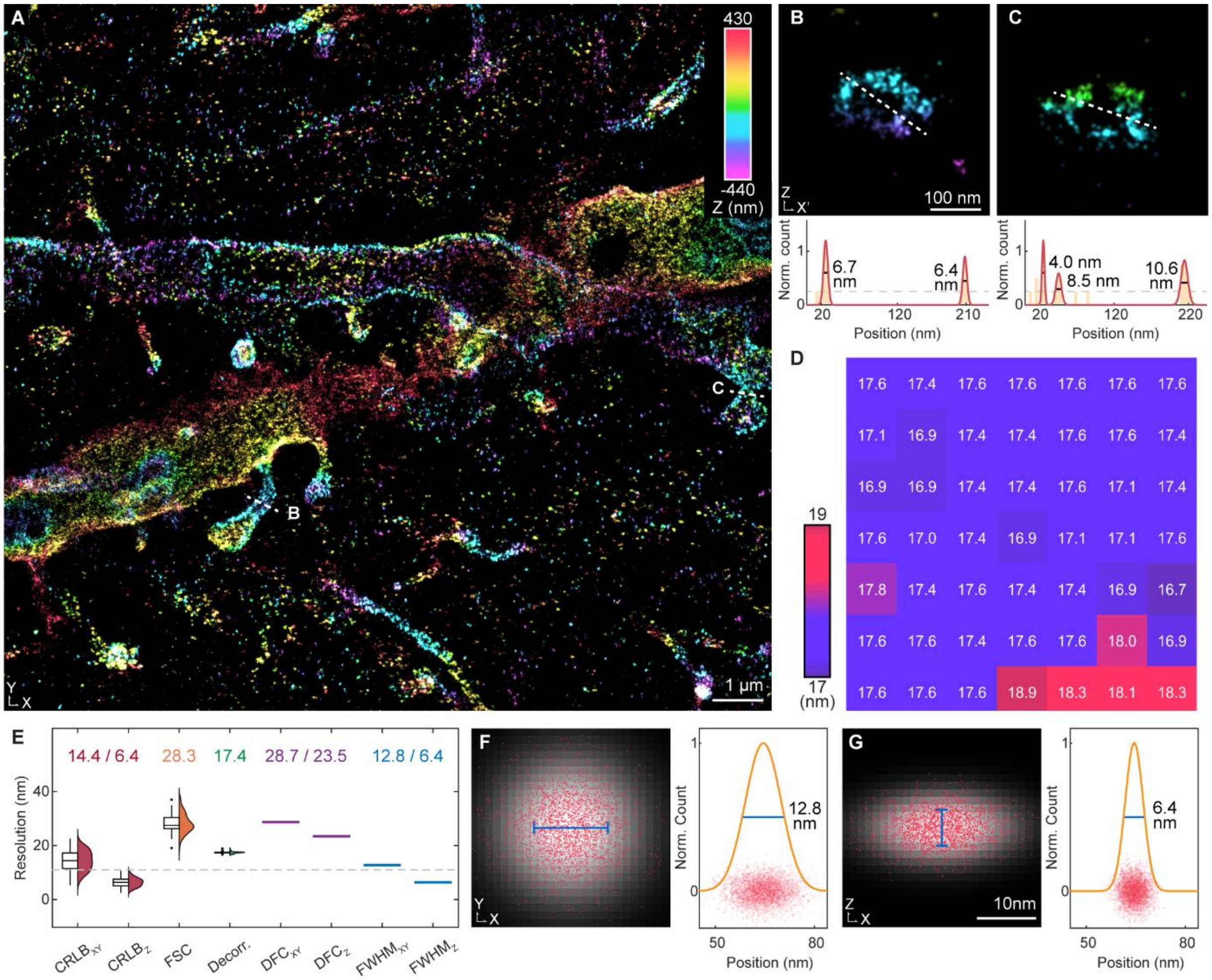


**Figure S7 Demonstration of 4Pi-BRAINSPOT in mouse brain slices. Related to Figure 5.**

(**A**) Reconstruction of AF647-labeled ChR2 in 50-μm thick mouse brain slices in Figure **6A**, color-coded according to the single-molecule Z positions.

(**B**&**C**) Vertical cross-sections (top) of targeting dendritic spines (white dashed lines) in (**A**). Normalized intensity profile (bottom) along white dashed lines. Yellow bar is localization histogram with 6-nm bins, red curve is fitted Gaussian functions to the histogram, gray dashed line is exclusion threshold, black solid line marks estimated feature FWHM, solid arrowhead indicates excluded isolated features, and dashed arrowhead indicates feature discontinuity.

(**D**) Heatmap of image decorrelation resolutions by segmenting 2D projection of (**A**) into 7x7 subregions.

(**E**) Comparison of resolution analysis using different methods, including lateral and axial CRLB-estimated resolutions ($CRLB_{XY}$ of 14.4±3.7 nm and $CRLB_Z$ of 6.4±1.6 nm, n=3244 localizations, red), FSC resolution (FSC of 28.3±3.6 nm, n=81 subregions, orange), imaging decorrelation resolution (Decor. of 17.4±0.4 nm, n=81 subregions, green), lateral and axial DFC resolutions ($DFC_{XY}$ of 28.7 nm and $DFC_Z$ of 23.5 nm, purple), and measured lateral and axial FWHMs of superposed single-molecule cluster ($FWHM_{XY}$ of 12.8 nm and $FWHM_Z$ of 6.4 nm, cyan). Box is upper and lower quartiles, central line is median, error bar is furthest data point excluding outliers, gray dashed line is estimated isotropic 3D CRLB-estimated resolution, and distribution is shown on the right side.

(**F**&**G**) Horizontal (**F**) and vertical (**G**) cross-sections (left) and normalized intensity profiles (right) of superposed single-molecule cluster. Red dot is single-molecule localization (n=3244 localizations), grayscale distribution is rendered cluster using 2D Gaussian blur with estimated FWHM, orange curve is fitted Gaussian function, and blue line marks estimated FWHM.

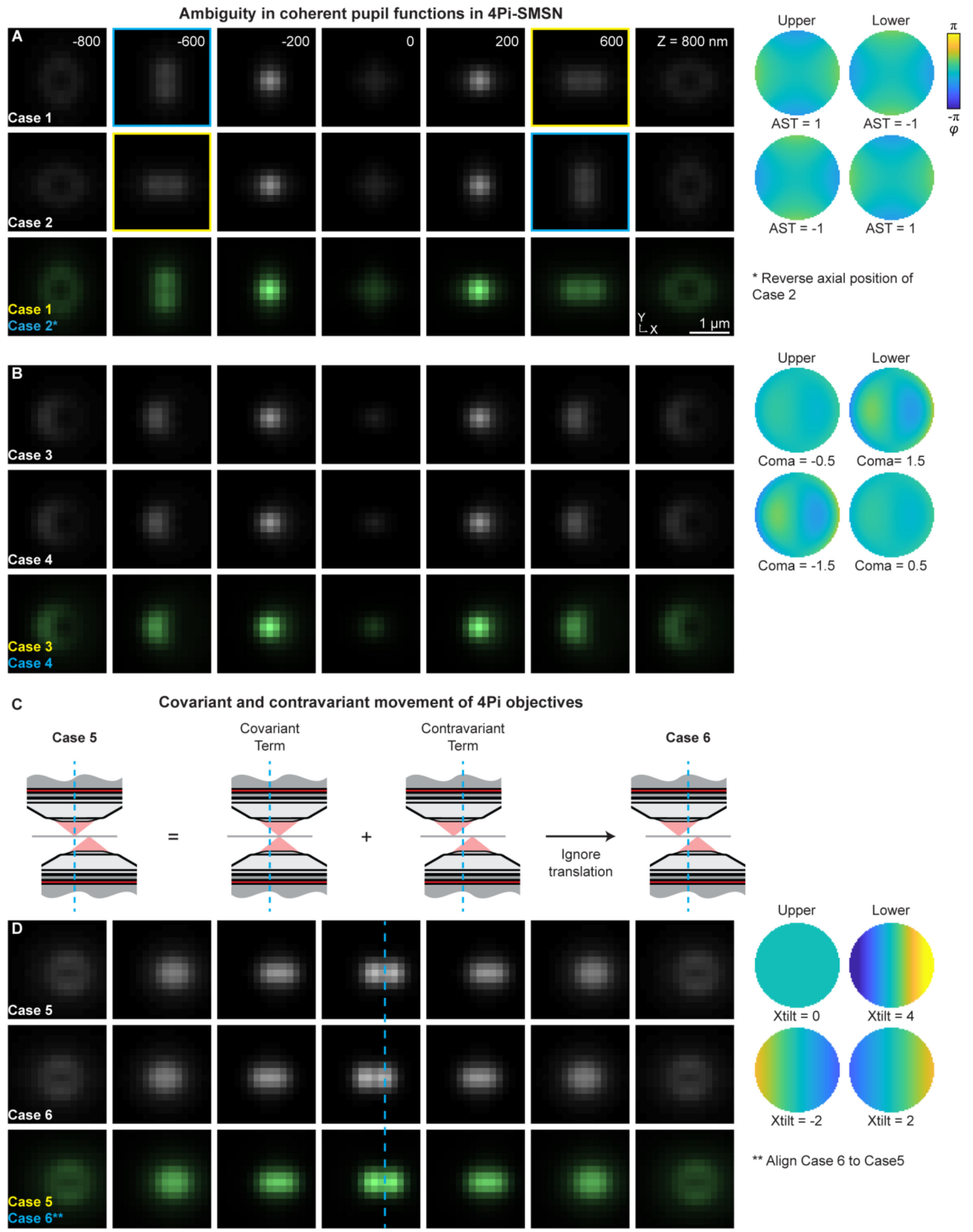


**Figure S8 Ambiguity in 4Pi-PSF configurations.**

(**A**) 4Pi-PSFs of channel p1 at z positions ranging from ±1 μm with vertical astigmatism variations of ±1 λ/2π (case 1, top row) and ∓1 λ/2π (case 2, middle row) in the upper and lower pupil functions, respectively.

Yellow and cyan boxes highlight identical PSFs with opposite Z positions in cases 1 and 2. Overlapping 4Pi-PSFs with axial symmetry ambiguity (bottom row) between case 1 (yellow) and case 2 with reversed sequence (cyan).

(**B**) 4Pi-PSFs of channel p1 at z positions ranging from ±1 μm with vertical coma aberrations of -0.5 and 1.5 $\lambda/2\pi$ (case 3, top row) and -1.5 and 0.5 $\lambda/2\pi$ (case 4, middle row) in the upper and lower pupil functions, respectively. Cases 3 and 4 show identical PSFs at all Z positions. Overlapping 4Pi-PSFs with indistinguishable ambiguity (bottom row) between case 3 (yellow) and case 4 (cyan).

(**C**) Effect of covariant and contravariant objective misalignment in 4Pi-SMSN.

(**D**) 4Pi-PSFs of channel p1 at z positions ranging from ±1 μm with vertical coma aberrations of -0 and 4 $\lambda/2\pi$ (case 5, top row) and -2 and 2 $\lambda/2\pi$ (case 6, middle row) in the upper and lower pupil functions, respectively. Cyan dashed line highlights the translation between identical PSF patterns in cases 5 and 6. Overlapping 4Pi-PSFs with indistinguishable translated ambiguity (bottom row) between case 5 (yellow) and case 6 with translation (cyan).

**Supplementary Methods**

**Resource table**

| REAGENT or RESOURCE | SOURCE | IDENTIFIER |
|---|---|---|
| Antibodies | | |
| Rabbit anti-Tom20 (FL-145) | Abcam | Cat# ab78547;<br>RRID: AB_2043078 |
| Mouse anti-alpha-Tubulin | Sigma-Aldrich | Cat#T5168;<br>RRID: AB_477579 |
| Sheep anti-NOGO-B (RTN4) | R&D systems | Cat#AF6034<br>RRID: AB_10573837 |
| Chicken anti-GFP | Abcam | Cat# ab13970<br>RRID: AB_300798 |
| Goat anti-rabbit IgG, Alexa Fluor 647 | Molecular Probes | Cat#A21236;<br>RRID: AB_141725 |
| Goat anti-mouse IgG, Alexa Fluor 647 | Abcam | Cat# ab150115;<br>RRID: AB_2687948 |
| Donkey anti-sheep, Alexa Fluor 647 | ThermoFisher Scientific | Cat# A21448;<br>RRID: AB_2535865 |
| Goat anti-chicken, Alexa Fluor 647 | ThermoFisher Scientific | Cat# A21449;<br>RRID: AB_2535866 |
| Chemicals, peptides, and recombinant proteins | | |
| PBS (1X), pH 7.4 | ThermoFisher Scientific | Cat# 10010049 |
| MES buffer | ThermoFisher Scientific | Cat# J60763-AP |
| NaCl | Sigma-Aldrich | Cat# S9888 |
| MgCl2 | ThermoFisher Scientific | Cat# AM9530G |
| EGTA | Sigma-Aldrich | Cat# E4378 |
| Sucrose | Sigma-Aldrich | Cat# 84097 |
| Poly-L-lysine solution | Sigma-Aldrich | Cat#P4707 |
| 2,2′-Thiodiethanol (TDE) | Sigma-Aldrich | Cat#166782 |
| Silicone dental glue | Picodent | Cat#Twinsil speed 22 |
| NaHCO3 | ThermoFisher Scientific | Cat#424270250 |
| Dulbecco's modified eagle medium (DMEM) | American Type Culture Collection | Cat#30-2002 |
| 10% Fetal bovine serum (FBS) | American Type Culture Collection | Cat#30-2020 |
| 1% penicillin-streptomycin | Gibco | Cat#15140122 |
| 0.2% saponin | Sigma-Aldrich | Cat#SAE0073 |
| 16% Paraformaldehyde aqueous solution (PFA in fixing cells and mouse brains) | Electron Microscopy Sciences | Cat#15710 |
| Paraformaldehyde Solution, 4% in PBS (PFA in fixing mouse forelimbs) | Fisher Scientific | Cat#J19943K2 |
| 8% Glutaraldehyde aqueous solution (GA) | Electron Microscopy Sciences | Cat#16019 |
| Triton X-100 | Sigma-Aldrich | Cat#X100 |
| Sodium borohydride (NaBH4) | Sigma-Aldrich | Cat#452882 |
| Bovine serum albumin (IgG-Free, Protease-Free) (BSA in labeling cells) | Jackson ImmunoResearch | Cat#001000162 |
| Bovine serum albumin (BSA in labeling mouse forelimbs) | Sigma-Aldrich | Cat#A9418 |
| 5% Donkey serum | Jackson ImmunoResearch | Cat#017000121 |

| | | |
|---|---|---|
| Tris (Base) | Avantor | Cat#JT4109 |
| 200 nm crimson beads | Invitrogen | N/A |
| Cysteamine hydrochloride (MEA) | Sigma-Aldrich | Cat#M6500 |
| 2-Mercaptoethanol (BME) | Sigma-Aldrich | Cat#M3148 |
| Cyclooctatetraene (COT) | Sigma-Aldrich | Cat#138924 |
| 3,4-Dihydroxybenzoic acid (PCA) | Sigma-Aldrich | Cat#37580 |
| Protocatechuate 3,4-dioxygenase (PCD) | Sigma-Aldrich | Cat#P8279 |
| D-(+)-glucose | ThermoFisher Scientific | Cat#A16828-36 |
| Ketamine | Akron | Cat#59399-114-10 |
| Xylazine | HVS | Cat#343750 |
| Urea | Fisher Scientific | Cat#U15-500 |
| D-sorbitol | Sigma-Aldrich | Cat#S1876-500G |
| glycerol | Sigma-Aldrich | Cat#G5516 |
| DMSO | Sigma-Aldrich | Cat#276855 |
| Experimental models: Cell lines | | |
| COS-7 | ATCC | Cat#CRL-1651<br>RRID: CVCL_0224 |
| U2OS | Cytion | Cat#300444<br>RRID: CVCL_B7FL |
| Experimental models: Organisms/strains | | |
| Mouse: B6.Cg-Tg(Thy1-COP4/EYFP)18Gfng/J | The Jackson Laboratory | Strain #:007612;<br>RRID: 007612 |
| Software and algorithms | | |
| MATLAB R2020a | MathWorks | www.mathworks.com/ |
| Dipimage 2.9 | Delft University of Technology | www.diplib.org/ |
| Blender 3.4 | Blender | www.blender.org/ |
| LabVIEW 2015 | National Instruments | www.ni.com/ |

**Sample preparation and data acquisition**

**1.1. Preparation and clearing of coverslip**

High-precision glass coverslips (25-mm diameter, CG15XH, Thorlabs) were cleaned through a series of steps to ensure their readiness for experimental use. The coverslips were first sonicated for 15 mins in a 1M KOH solution using an ultrasonic cleaner (M2800H, Branson). This was followed by a 15-min sonication in double-distilled water and another 15-min sonication in 70% ethanol. Between each sonication step, the coverslips were rinsed with double-distilled water to remove any residual liquid from their surfaces. After the sonication steps, the coverslips were placed in a 110 °C oven (89508-424, Avantor) and heated until they were completely dry.

**1.2. Sample preparation of fluorescent bead on coverslip**

The cleaned coverslips were coated with poly-L-lysine solution (P4707, Sigma-Aldrich) by immersing them for 30 mins at room temperature (RT) and rinsed once with double-distilled water. The coverslips were then incubated with 100 µL of a 1:$10^6$ diluted solution of 100-nm crimson beads (100 nm crimson beads, Invitrogen) in double-distilled water for 30 mins at RT, followed by another rinse with double-distilled water. The samples were mounted by adding 50 µL of 97% 2,2'-thiodiethanol (TDE, 166782, Sigma-Aldrich) in double-distilled water between the treated coverslips and cleared coverslips. The sandwiched samples were sealed using silicone dental glue (Twinsil speed 22, Picodent) and left to dry at RT for 30 mins prior to imaging.

**1.3. Sample preparation of fluorescent dye on coverslip**

The cleaned coverslips were immersed with poly-L-lysine solution (P4707, Sigma-Aldrich) for 30 mins at room temperature (RT) and rinsed once with double-distilled water. The coverslips were incubated with 100 µL of a 1:$10^3$ diluted Alexa 647 (A21236, Molecular Probes) in 0.1M NaHCO3 for 30 mins at RT, followed by two rinses with 0.1M NaHCO3. The samples were mounted by adding 50 µL of a mixture containing 2.5 mM 3,4-dihydroxybenzoic acid (PCA, 37580, Sigma-Aldrich), and 50 nM protocatechuate 3,4-dioxygenase (PCD, P8279-25UN, Sigma-Aldrich) in dSTORM buffer (10% wt/vol glucose in 50 mM Tris, 50 mM NaCl, pH 8.0) between the treated coverslips and the other cleared coverslips. The sandwiched samples were sealed using silicone dental glue and left to dry at RT for 30 mins prior to imaging.

**1.4. Cell culture**

COS-7 cells (CRL-1651, ATCC) were grown on clean coverslips placed in six-well plates. Before adding cells, the six-well plates were exposed to UV light for 1 hr. The cells were cultured in Dulbecco's modified eagle medium (DMEM, 30-2002, ATCC) supplemented with 10% v/v fetal bovine serum (FBS, A5670701,

Gibco) and 100 U/mL penicillin–streptomycin (P-S, 15070063, Gibco). The culture conditions were maintained at 37 °C with 5% $CO_2$ in a humidified incubator until the confluence of the cells reached approximately 60%. The cells were grown on coverslips for a duration ranging from 12 to 24 hours before fixation.

U2OS Cells (300444, Cytion) were grown on clean coverslips placed in six-well plates. Before adding cells, the six-well plates were exposed to UV light for 1 hr. The cells were cultured in McCoy's 5A (16600, Gibco) supplemented with 10% v/v fetal bovine serum (FBS, A5670701, Gibco) and 100 U/mL penicillin-streptomycin. The culture conditions were maintained at 37 °C with 5% $CO_2$ in a humidified incubator until the confluence of the cells reached approximately 60%. The cells were grown on coverslips for a duration ranging from 12 to 24 hours before fixation.

#### 1.5. Fixation and immunofluorescence labeling of cells

The cell fixations and immunolabeling were tailored for different cellular structures. For immunofluorescence-labeled TOM20, cells were fixed with pre-warmed (37 °C) 4% paraformaldehyde (PFA, 1:4-diluted 16% paraformaldehyde solution, 15710, Electron Microscopy Sciences) in 1xPBS at RT for 15 mins with gentle rocking, followed by rinsing twice with 1× PBS. Autofluorescence quenching was performed with fresh 0.1% sodium borohydride ($NaBH_4$, 452882, Sigma-Aldrich) in 1× PBS at RT for 7 minutes with gentle rocking, followed by three rinses with 1× PBS. Cell permeabilization was carried out using block buffer (3% bovine serum albumin (BSA) and 0.2% Triton X-100 in 1× PBS) at RT for 30 mins with gentle rocking, after which the solution was aspirated. The cells were incubated with primary antibodies (1:2000-diluted rabbit anti-TOM20 primary antibody (ab78547, Abcam) in antibody dilution buffer (1% BSA and 0.2% Triton X-100 in 1× PBS)) at 4 °C overnight and rinsed three times for 5 mins with wash buffer (0.05% Triton X-100 in 1× PBS). The cells were then incubated with secondary antibodies (1:500-diluted goat anti-rabbit AF647 secondary antibody (A21236, Life Technologies) in antibody dilution buffer) at 4 °C for 4 hours and rinsed three times for 5 mins with wash buffer. Post-fixation was performed with the same buffer as fixation at RT for 10 mins with gentle rocking, followed by three rinses with 1× PBS. The cells were stored in 1× PBS at 4 °C until imaging.

For immunofluorescence-labeled α-tubulin, cells were pre-extracted with pre-warmed 0.2% saponin (SAE0073, Sigma-Aldrich) in cytoskeleton buffer (CBS, 10 mM MES pH 6.0, 138 mM NaCl, 3 mM MgCl2, 2 mM EGTA, 320 mM sucrose) at RT for 1 min with gentle rocking, after which the solution was aspirated. Subsequently, the cells were fixed with pre-warmed 3% PFA and 0.1% glutaraldehyde (GA, 1:80-diluted 8% glutaraldehyde solution, 16019, Electron Microscopy Sciences) in CBS at RT for 15 mins with gentle rocking, followed by rinsing twice with 1× PBS. Cell permeabilization was carried out using block buffer at RT for 30 mins with gentle rocking, after which the solution was aspirated. The cells were incubated with primary antibodies (1:500-diluted mouse anti-α-tubulin primary antibody (T5168, Sigma-Aldrich) in antibody dilution buffer) at 4 °C overnight and rinsed three times for 5 mins with wash buffer (0.05% Triton X-100 in 1× PBS).

The cells were then incubated with secondary antibodies (1:500-diluted goat anti-mouse AF647 secondary antibody (ab150115, Abcam) in antibody dilution buffer) at RT for 30 mins and rinsed three times for 5 mins with wash buffer. Post-fixation was performed with the same buffer as fixation at RT for 10 mins with gentle rocking, followed by three rinses with 1× PBS. The cells were stored in 1× PBS at 4 °C until imaging.

For immunofluorescence-labeled Rtn4, cells were fixed with pre-warmed 3% PFA and 0.5% GA in 1xPBS at RT for 15 mins with gentle rocking and rinsed twice with 1× PBS. Autofluorescence quenching was performed with fresh 0.1% sodium borohydride in 1× PBS at RT for 7 minutes with gentle rocking, followed by three rinses with 1× PBS. Cell permeabilization was carried out using 5% donkey serum (017000121, Jackson Immunoresearch) and 0.2% Triton X-100 in 1× PBS at RT for 30 mins with gentle rocking, after which the solution was aspirated. The cells were incubated with primary antibodies (1:200-diluted sheep anti-Rtn4b primary antibody (AF6034, R&D systems) in 5% donkey serum) at 4 °C overnight and rinsed three times for 5 mins with wash buffer. The cells were then incubated with secondary antibodies (1:400-diluted donkey anti-sheep AF647 secondary antibody (A21448, ThermoFisher Scientific) in block buffer) at RT for 10 mins and rinsed three times for 5 mins with wash buffer. Post-fixation was performed with the same buffer as fixation at RT for 10 mins with gentle rocking, followed by three rinses with 1× PBS. The cells were stored in 1× PBS at 4 °C until imaging.

#### 1.6. Sample mounting of cells and preparation of imaging buffer

Right before imaging acquisition, 1 μL of 1:$10^6$-diluted 200-nm crimson beads (200 nm crimson beads, Invitrogen) in double-distilled water were added around the center region of the coverslips with labeled cells on top. The samples were mounted with 70 μL of a freshly prepared imaging buffer consisting of 10 mM 2-mercaptoethylamine (MEA, M9768, Sigma-Aldrich), 50 mM 2-hydroxyethylmercaptan (BME, 63689, Sigma-Aldrich), 2 mM cyclooctatetraene (COT, 138924, Sigma-Aldrich), 2.5 mM PCA, and 50 nM PCD in dSTORM buffer. Another coverslip was placed on top to form a sandwiched sample. The samples were then sealed using silicone dental glue and left to dry at RT for 30 mins prior to imaging.

For the preparation of cell-on-top-coverslip samples, cleaned coverslips were initially immersed in poly-L-lysine solution at RT for 30 mins and subsequently rinsed once with double-distilled water. These coverslips were then incubated with 100 μL of 1:$10^6$-diluted 200-nm crimson beads in double-distilled water at RT for 30 mins and rinsed again with double-distilled water, serving as the bottom coverslips in the sample sandwiches. For the top coverslips in the sample sandwiches, coverslips with labeled cells on top were treated with 1 μL of 1:$10^6$-diluted 200-nm crimson beads in double-distilled water around the center region. The treated top and bottom coverslips were then assembled with 70 μL of freshly prepared imaging buffer, sealed using silicone dental glue, and left to dry at RT for 30 mins prior to imaging.

### 1.7. Immunofluorescence labeling of brain sections

The 3.5-month-old transgenic mice (B6.Cg-Tg(Thy1-COP4/EYFP)18Gfng/J, The Jackson Laboratory) were anesthetized with an intraperitoneal injection of a mixture containing 90 mg/kg ketamine (59399-114-10, Akron) and 10 mg/kg xylazine (343750, HVS). The mice were transcardially perfused with 1xPBS, followed by 4% PFA to pre-fix the brains. The brains were carefully excavated from the skulls and immersed in 4% PFA overnight at 4 °C for further fixation. The following day, fixed brain samples were cut into 50-μm coronal sections using a vibratome (1000 Plus, Vibratome). The sections were stored at 4 °C in 1x PBS until labeling.

For immunofluorescence-labeled Thy1+ pyramidal cells expressing ChR2-EYFP in mouse brains, the sections were rinsed with 0.1% Triton X-100 in 1× PBS three times for 15 mins with gentle rocking. They were then blocked with blocking buffer (5% BSA in 1× PBS) for 1.5 hours with gentle rocking. The sections were incubated with 1:1,000-diluted chicken anti-GFP antibody (ab13970, Abcam) in blocking buffer at 4 °C overnight and then rinsed three times for 15 mins with wash buffer (0.1% Triton X-100 in 1× PBS). Subsequently, the sections were incubated with 1:600-diluted goat anti-chicken AF647 secondary antibody (A21449, Invitrogen) in wash buffer at RT for 2 hours with gentle rocking. The sections were rinsed three times for 15 mins with wash buffer and stored in 1× PBS at 4 °C until imaging.

### 1.8. Tissue clearing for single-molecule localization microscopy

Tissue clearing is critical for high-resolution 4Pi-SMSN imaging, where light scattering and absorption degrade interferometric contrast, fluorescent signal strength, and background suppression. While tissue imaging protocols can largely follow those established for cultured cells once sufficient optical transparency is achieved, the inherent optical inhomogeneity of tissues limits the retrievable information from tissue imaging. To address this, tissue clearing methods have been developed to enhance transparency through lipid removal, decolorization, and decalcification, such as SeeDB, Scale, and iDISCO [1-6]. Established methods have demonstrated their ability to render brain tissue transparent while preserving structural integrity, enabling imaging depths from <20 μm to >150 μm with minimal sample distortion. However, not all methods are compatible with 4Pi-SMSN. Certain reagents commonly used in clearing protocols can disrupt the thiol-based imaging buffer required for single-molecule localization. In our experiment, tissue clearing methods incorporating sodium dodecyl sulfate, acrylamide hydrogel, dichloromethane, and dibenzyl ether affected fluorophore stability and single-molecule blinking, making them unsuitable for 4Pi-SMLM. In contrast, the fast optical clearing method (FOCM) [7] provided tissue transparency while maintaining acceptable blinking performance when integrated with thiol-based imaging buffers. The tissue clearing was performed by incubating brain section with 200 μL FOCM reagent (30% wt/vol urea (U15-500, Fisher Scientific), 20% wt/vol D-sorbitol (S1876-500G, Sigma-Aldrich), and 5% wt/vol glycerol (G5516, Sigma-Aldrich) in DMSO (276855, Sigma-Aldrich)) at RT for 1-2 mins.

#### 1.9. Sample mounting of brain sections and tissue clearing

Right before imaging acquisition, the labeled brain sections were placed on cleaned coverslips, with the visual cortex regions positioned at the center of the coverslips. Tissue clearing was then performed and rinsed once with 1xPBS. 1 µL of 1:$10^6$-diluted 200-nm crimson beads in double-distilled water was added around the outer edge of the visual cortex. The samples were then assembled with 70 µL of freshly prepared imaging buffer, sealed using silicone dental glue, and left to dry at RT for 30 mins prior to imaging.

For the preparation of bead with unlabeled brain section samples, cleaned coverslips were initially immersed in poly-L-lysine solution at RT for 30 mins and subsequently rinsed once with double-distilled water. These coverslips were then incubated with 100 µL of 1:$10^6$-diluted 200-nm crimson beads in double-distilled water at RT for 30 mins and rinsed again with double-distilled water. The unlabeled brain sections were placed on the center of the coverslips, serving as the bottom coverslips in the sample sandwiches. For the top coverslips in the sample sandwiches, coverslips with labeled cells on top were treated with 1 µL of 1:$10^6$-diluted 200-nm crimson beads in double-distilled water around the center region. The treated top and bottom coverslips were then assembled with 70 µL of 1x PBS, sealed using silicone dental glue, and left to dry at RT for 30 mins prior to imaging.

#### 1.10. Imaging acquisition of fluorescent beads and system calibration

The bead sample was illuminated with a modest laser power of 30 W/cm$^2$ for calibration and imaging acquisition. The alignment process began with a preliminary alignment of the upper and lower objectives to coincide with fluorescent patterns acquired from them. Next, the upper and lower DMs were meticulously calibrated to introduce vertical astigmatism with amplitudes of ±1 $\lambda/2\pi$, preventing axial ambiguity, and minimizing systematic aberrations. To estimate the aberrations, beads were scanned within an axial range of ±1 µm, and acquired using upper and lower objectives, respectively. The step size for this scan was 100 nm, and each position was recorded for 5 frames with an acquisition time of 100 ms. The resulting bead measurements were subjected to a phase retrieval algorithm to calculate the respective pupil functions and their corresponding Zernike decomposition [8,9]. To compensate for systematic aberrations, voltage maps corresponding to the opposite Zernike modes (Wyant order, up to 2nd-order aberrations) were applied to the DMs for calibration. To optimize the interferometric contrast in the coherent fluorescent detection, the tower mechanical system was scanned to measure the interferometric intensity profile at the bead's center under different cavity phases. The tower mechanical system was then positioned at the position where the maximal intensity modulation was observed.

In the experiment assessing the practical performance of the *in-situ* 4Pi-PSF modeling, an interferometric bead measurement was conducted. The beads were scanned within an axial range of ±1 µm. The step size was set at 100 nm, and each position was recorded for 50 frames, with an acquisition time of 20 ms. In addition to the non-aberrated measurements, 7 Zernike modes (Wyant order, ranging from vertical astigmatism to vertical trefoil) were introduced to the upper and lower DMs with amplitudes of ±1 λ/2π, creating aberrated interferometric patterns.

#### 1.11. Imaging acquisition of biological samples and measurement of sample thickness

The biological sample was illuminated with a modest laser power of 30 W/cm$^2$ for searching region of interest, after which the system was calibrated employing beads on the coverslip (Method 1.10). For the construction of the *in-vitro* cubic-spine model, beads were scanned within an axial range of ±1 μm to capture the interferometric patterns [10]. The step size for this scan was set at 20 nm, and each position was recorded for 5 frames with an acquisition time of 100 ms. In constructing the *in-vitro* phase-retrieval model [11], individual pupil functions of both objectives were collected, using the same procedure in fluorescent bead imaging acquisition (Method 1.10), besides interferometric bead measurement.

Afterward, the samples were positioned to center the targeted structures, followed by exposure to a laser with a power of 12 kW/cm² for 1 min, transferring fluorophores into dark states and bleached autofluorescence. Subsequently, the laser power was reduced to 7 kW/cm² for imaging acquisition. For each FOV, 80,000-100,000 frames were collected with an acquisition time of 20 ms. Each frame was then cropped to yield four-channel detections. In the experiment evaluating the dynamic update capability of 4Pi *in-situ* PSF modeling, we collected 20 sequential datasets, each comprising 2,000 frames, following the above configuration. Prior to each collection window, random aberrations were introduced to the deformable mirrors in the upper and lower arms. These aberrations were generated across 10 Zernike modes (Wyant order, ranging from vertical tilt to vertical trefoil), with random amplitudes between ±1 λ/2π. To measure the sample thickness at the targeted position, the sample was incrementally moved through the focal plane. The distance between the bottom and top coverslip surfaces, identified by the presence of random dust particles on the top surface, was recorded as the sample thickness.

### Systematic configuration and simulated method

#### 2.1 4Pi single molecule localization microscopy

The 4Pi single molecule imaging system was assembled based on a previously published schematic design[11-13], incorporating two 100x/1.4-NA oil-immersion objectives (UPLSAPO 100XO, Olympus) for coherent interferometric single-molecule detection. The emission path was configured to collect emissions from both objectives, which were divided into P- and S-polarized orientations with equal amplitudes using quarter-wave plates (AQWP05M-600-UM-SP, Thorlabs). The emissions were aligned along four coinciding paths using a combination of nonpolarized and polarized beam splitters (NT47-009 and NT49-002, Edmund Optics), and directed onto distinct regions of an sCMOS camera (Orca-Flash4.0v2, Hamamatsu) for simultaneous 4-channel acquisition. A telecentric optical design was employed, resulting in a final magnification of 50x and an effective pixel size of 129 nm, allowing for 4Pi interferometric imaging across a 20-μm field of view (FOV).

To reduce background signals in fluorescent acquisition, quad-band dichroic and bandpass filters (FF01-446/523/600/677-25, Di01-R405/488/561/635-17.5×24, Semrock) were positioned between the quarter-wave plates and the nonpolarized beam splitter. This arrangement separated the emission from the excitation. Aberrations in two interferometric arms were individually controlled using two deformable mirrors (DM) (Multi-5.5, Boston Micromachines), equipped with 140 actuators, placed at the conjugate pupil planes. To control phase delay and dispersion across the four optical paths, two customized Babinet-Soleil compensators were placed between the DMs and the nonpolarized beam splitter. The thickness of the compensators was optimized to achieve phase delays of 0, π/2, π, and 3π/2 with minimized dispersions between the upper and lower arms for four channels.

In excitation, a 642-nm continuous-wave laser (2RU-VFL-P-2000-642-B1R, MPB Communications) was used, whose power was controlled by an acousto-optic tunable module (AOTFnC-400.650-TN, AA Opto-electronic). The laser light was guided through a polarization-maintaining single-mode fiber (PM-S405-XP, Thorlabs) and focused on the back focal plane of the lower objective for Köhler illumination, generating an illumination area of 27 μm. A mirror positioned at the conjugate imaging plane allowed for switching tilted angles of illumination between 0° (Epi mode) and 54° (HILO mode).

For alignment of the interferometric cavity, the upper and lower objectives were mounted on piezo nano-positioning motors (N-664.3A, P-612.2SL, PI) for axial and lateral adjustments, respectively. The sample holder was arranged on a combination of piezo positioning motors (P-541.XYZ, M-227.10, M-686.D64, PI) for nanoscale sample positioning within a microscale range. A tower mechanical system (LS-50, ASI) facilitated microscale vertical positioning of the objective and sample positioning modules. The nonpolarized beam splitter was mounted on a rotation goniometric motor (M-GON40-U, M-RS65, TRA12CC, Newport) for fine-tuning the spatial coincidence of upper and lower fluorescence. The microscope setup was synchronized using a custom LabVIEW program.

#### 2.2 Aberration control and calibration of deformable mirrors

The DM calibration followed established methodologies, where DMs are controlled by an orthogonal set of 32 mirror deformation modes (DM modes, corresponding to Zernike modes up to 4$^{th}$-order spherical aberration) [13-16]. To measure the response of DM modes, 100-nm fluorescent beads were scanned within an axial range of ±1 μm to capture the single-molecule patterns corresponding to DM modes with amplitudes of ±1 λ/2π. The step size for this scan was 400 nm, and each position was recorded for 3 frames with an acquisition time of 100 ms. The resulting bead measurements were subjected to a phase retrieval algorithm to calculate the respective pupil functions for certain DM modes [8,9].

To solve mismatched orientations of the two DMs, we decomposed the DM modes of each DM into Zernike modes with the same orientation. The pupil function of the m$^{th}$ DM mode $\Phi_m$ was decomposed into a series of Zernike modes, excluding Piston, X-tilt, Y-tilt, and defocus aberration, which was expressed as

$$\Phi_m(k_x, k_y) \cong \sum_{n=5}^{64} c_n Z_n(k_x, k_y) \tag{1}$$

where $c_n$ and $Z_n(k_x, k_y)$ are coefficient and aberration phase of the $n^{th}$ Zernike mode.

By solving this equation with least-square minimization, each Zernike mode can be expressed as a linear combination of DM modes. To mitigate the impact of residual aberrations, the difference between the retrieved amplitudes at amplitudes of ±1 λ/2π was calculated and divided by 2. To assess the performance of DM control, we retrieved pupil functions from fluorescent bead measurements by applying 17 Zernike aberrations (following Wyant order, from vertical astigmatism to oblique quadrafoil) with amplitudes of ±1 λ/2π. The beads were scanned over an axial range of ±1 μm, with a step size of 100 nm, and each position was recorded for 5 frames with an acquisition time of 100 ms. The retrieved pupil functions showed an error of less than 0.4 λ/2π from the applied Zernike modes.

#### 2.3 Characterization of sCMOS camera

The characterization of the sCMOS camera, including parameters such as offset, variance, and gain for each pixel, was determined using established methodologies [17]. For pixel $i$, the relationship between input photon $X_i$ and the readout value $Y_i$ can be described as

$$Y_i = g_i \cdot Poisson(X_i) + Gaussian(o_i, \sigma_i^2) \tag{2}$$

where $g_i$ is the gain of the pixel $i$, $o_i$ and $\sigma_i^2$ are Gaussian mean and variance of the pixel $i$. $Poisson(x)$ represents Poisson random process with mean of $x$, and $Gaussian(o, \sigma^2)$ represents Gaussian random process with mean of $o$ and variance of $\sigma^2$.

The Gaussian mean $o_i$ and variance $\sigma_i^2$ can be estimated with illumination photon $x_i = 0$ as

$$\widehat{o_i} = E_{t \in T}[y_i^t | x_i^t = 0] \tag{3}$$

$$\widehat{\sigma_i}^2 = Var_{t \in T}[y_i^t | x_i^t = 0] \tag{4}$$

where $x_i^m$ and $y_i^m$ are the input photon and readout value of pixel $i$ at frame $t$, $t \in T$ encompasses all valid frames. The gain factor $g_i$ can then be estimated under different illumination photon levels $x_i = s_n$ by solving the least-square minimization problem

$$\widehat{g_i} = \arg\min \|B - g_i A\|^2 \tag{5}$$

$$\text{where } A = \begin{bmatrix} E_{t\in T}(y_i^m | x_i^m = s_1) - \widehat{o_i} \\ E_{t\in T}(y_i^m | x_i^m = s_2) - \widehat{o_i} \\ \vdots \\ E_{t\in T}(y_i^m | x_i^m = s_n) - \widehat{o_i} \end{bmatrix}, \quad B = \begin{bmatrix} Var_{t\in T}(y_i^m | x_i^m = s_1) - \widehat{\sigma_i}^2 \\ Var_{t\in T}(y_i^m | x_i^m = s_2) - \widehat{\sigma_i}^2 \\ \vdots \\ Var_{t\in T}(y_i^m | x_i^m = s_n) - \widehat{\sigma_i}^2 \end{bmatrix}$$

The estimation of this overdetermined system can be derived by Moore–Penrose pseudo-inverse as

$$g_i = (AA^T)^{-1}AB \tag{6}$$

To assess the characterization experimentally dye-on-coverslip samples were imaged under illumination levels ranging from 0 to 3 kW/cm$^2$ for 5 trials. For each trial, 2,000 frames were recorded with an acquisition time of 20 ms. The $o_i$, $\sigma_i^2$, and $g_i$ were then estimated using Eq. 3-6.

### 2.4 Coherence emission of 4Pi-SMSN

The 4Pi coherent emission can be described through the superposition of fluorescent wave fields collected from opposing objectives with the coincided focal plane, forming constructive or destructive interferometric patterns. The pupil functions of the system represent the amplitude of fluorescent wave fields at the conjugated pupil plane can be expressed as

$$h(k_x, k_y) = \begin{cases} a(k_x, k_y) \cdot e^{i\varphi(k_x, k_y)}, & \text{if } {k_x}^2 + {k_y}^2 \le \left(\frac{NA}{\lambda}\right)^2 \\ 0, & \text{if } {k_x}^2 + {k_y}^2 > \left(\frac{NA}{\lambda}\right)^2 \end{cases} \tag{7}$$

where $a(k_x, k_y)$ and $\varphi(k_x, k_y)$ are the magnitude and phase of the electric field at the pupil plane, respectively, $k_x$ and $k_y$ are wave vectors along X and Y directions, $NA$ is the numerical aperture of the objectives and $\lambda$ is the fluorescent wavelength in air. The phase of pupil function $\varphi(k_x, k_y)$, which include sample- and system-induced interferometric aberrations, can be decomposed into a series of Zernike modes as

$$\varphi(k_x, k_y) = \sum_{n=1}^{N} c_n Z_n(k_x, k_y), \tag{8}$$

where $Z_n(k_x, k_y)$ is the $n^{th}$ Zernike mode, $c_n$ is corresponding amplitude, and $N$ is the total number of Zernike modes. The complexity of interferometric electric fields in 4Pi emission, resulting from superposition of coherent pupil functions from the upper and lower objectives, $h_U$ and $h_L$, can be described by

$$\begin{aligned} h_{s1}(k_x, k_y, z) &= h_U(k_x, k_y)D(-z)e^{i\pi} + I_t h_L(k_x, k_y)D(z)e^{i(\varphi_{sp}+\varphi_0)} \\ h_{s2}(k_x, k_y, z) &= h_U(k_x, k_y)D(-z) + I_t h_L(k_x, k_y)D(z)e^{i(\varphi_{sp}+\varphi_0)} \\ h_{p1}(k_x, k_y, z) &= h_U(k_x, k_y)D(-z) + I_t h_L(k_x, k_y)D(z)e^{i(\pi+\varphi_0)} \end{aligned} \tag{9}$$

$$h_{p2}(k_x, k_y, z) = h_U(k_x, k_y)D(-z) + I_t h_L(k_x, k_y)D(z)e^{i\varphi_0}$$

where $D(z) = e^{i2\pi k_z(k_x,k_y)z}$ is defocusing term of wave fields, $k_z(k_x, k_y) = \sqrt{\left(\frac{n}{\lambda}\right)^2 - {k_x}^2 - {k_y}^2}$ is wave vector along Z direction, $n$ is the refractive index of the medium, $\varphi_{sp}$ is the phase difference between s- and p-polarized fluorescence, $\varphi_0$ is cavity phase between upper and lower paths when the emitter is at the coincided focal plane, and $I_t$ is the transmission efficiency ratio between two optical paths.

### 2.5 Interferometric modeling of 4Pi-PSF

The 4Pi-PSF can be represented as Fourier transform of interferometric electric fields. According to scalar diffraction theory, the intensity $\mu_{Im}$ and amplitude $H_m$ of a coherent 4Pi-PSF for channel $m \in (s1, s2, p1, p2)$ can be described as

$$\mu_{Im}(x, y, z) = |H_m(x, y, z)|^2, m \in (s1, s2, p1, p2) \quad (10)$$

$$H_m(x, y, z) = A_m(x, y, z) \cdot e^{i\Phi_m(x,y,z)} = \mathcal{F}\left[h_m(k_x, k_y, z)\right] \quad (11)$$

where $A_m(x, y, z)$ and $\Phi_m(x, y, z)$ are magnitude and phase of $H_m(x, y, z)$, $\mathcal{F}$ denotes the Fourier transform operator, and $h_m(k_x, k_y, z)$ is the interferometric wavefields for channel $m$ in Eq. 9. In practical implementations, the interference may exhibit partial coherence, leading to reduced fringe contrast due to any incoherent fluorescence. To account for this, the partially coherent 4Pi-PSF $\mu_0$ is expressed as a linear combination of coherent intensity $\mu_I$ and incoherent intensity $\mu_C$, which can be described as

$$\mu_{0m}(x, y, z) = a\mu_{Im}(x, y, z) + (1 - a)\mu_C(x, y, z) \quad (12)$$

$$\mu_C(x, y, z) = \left|\mathcal{F}[h_U(k_x, k_y)D(-z)]\right|^2 + \left|\mathcal{F}[I_t h_L(k_x, k_y)D(z)]\right|^2 \quad (13)$$

where $a$ is wavelength-dependent coherence factor between 0 and 1.

### 2.6 Simulation of 4Pi-PSF and determination of hyperparameters

The 4Pi-PSF simulation began by initiating pupil functions of $h_U$ and $h_L$, which incorporated optical aberrations by setting Zernike modes. The partially coherent 4Pi-PSF $\mu_{0m}$ for channel $m$ was then generated based on Eq. 9-12 with hyperparameters of $NA$, $\lambda$, $n$, $\varphi_{sp}$, $I_t$, and $a$. To account for noise processing, multiplication of $\mu_{0m}$ and photon counts serve as the input to model the readout values of the sCMOS camera in Eq. 2.

To simulate practical imaging conditions for characterizing our algorithmic performance, we determined the initiating hyperparameters from real data. The emission wavelength $\lambda$ was estimated from acquisition of crimson beads with an emission spectrum similar to AF647 and optimized by minimizing step size estimation errors. The refractive indices $n$ of the imaging buffer and immersion oil were measured to be

1.352 and 1.516, respectively, using an Abbe refractometer (334610, Thermo Spectronic). The transmission efficiency $I_t$ was measured by single-molecule photon counts collected from the upper and lower objectives.

To estimate the coherence factor $a$, we employed published configuration based on Gaussian-weighted moment operators [12]. The process begins with aligning 4-channel interferometric patterns (Method 3.2). Gaussian-weighted moment operators $m(z)$ at z position are calculated by

$$m_m(z) = \iint D_m(x, y, z)\sqrt{x^2 + y^2} e^{-\frac{x^2+y^2}{2\sigma^2}} dxdy \tag{14}$$

where $D_m(x, y, z)$ is the acquired photons for channel $m$, and $\sigma$ is predetermined root-mean-square (RMS) Gaussian width of 77 nm. According to 4Pi emission, the phase delays between channels with the same polarization are π rad. This implies that locally maximal and minimal values of two moment operators with the same polarization will occur at the same axial positions, respectively, where the coherence factor $a$ can be estimated by calculating the maximal fringe contrast of

$$a = \frac{a_p + a_s}{2}$$
$$, where\ a_k = \max \frac{|m_{k1}(z) - m_{k2}(z)|}{m_{k1}(z) + m_{k2}(z)}, k \in (s, p) \tag{15}$$

To estimate the phase difference between s- and p-polarized fluorescence $\varphi_{sp}$, we further express the moment operators $M(z)$ as linear combinations of coherent terms and incoherent terms as

$$M_{m'}(z) = A_s(z)[d_{m'} \pm b_{m'} \cos(k_z z + \varphi_s)], m' \in (s1, s2)$$
$$M_{m''}(z) = A_p(z)\left[d_{m''} \pm b_{m''} \sin\left(k_z z + \varphi_p\right)\right], m'' \in (p1, p2) \tag{16}$$

where $A_s(z) = \sqrt{m_{s1}(z)^2 + m_{s2}(z)^2}$ and $A_p(z) = \sqrt{m_{p1}(z)^2 + m_{p2}(z)^2}$ are the acquired modulation envelopes of s- and p-polarization, $d_{m'}$ and $b_{m'}$ are the incoherent and coherent terms of the operator modulation, $k_z$ is modulation frequency, and $\varphi_s$ and $\varphi_p$ is phase delay of s- and p-polarization. The phase difference $\varphi_{sp} = \varphi_s - \varphi_s$ was estimated from normalized operator modulation using the 'fminsearch' function in MATLAB by solving the minimization problem as

$$\hat{\theta} = \arg\min \sum_m \left(\frac{M_m(\theta|z)}{A_k(z)} - \frac{m_m(z)}{A_k(z)}\right)^2, \theta \in \left(\varphi_s, \varphi_p\right) \tag{17}$$

which was ranging from -1.5 to -1.7 rad in our setup.

In conducting simulations of randomly aberrated 4Pi-PSF, we set the amplitude of vertical astigmatism to -1.5 and +1.5 λ/2π for the upper and lower pupils as prior knowledge. The amplitudes of other Zernike modes, ranging from oblique astigmatism to tertiary spherical aberration, were randomly sampled within a range of ±1 λ/2π for 30 trials. In each trial, we generated 2,000 interferometric patterns at random axial positions ranging from ±800 nm with random lateral offsets of ±250 nm from the center within a sub-region size of 40 pixels. The photon count for each emission pattern was set to 2,000 and the background count per pixel was set to 5.

In conducting simulations of cavity phase estimation, the above aberration configuration was used to generate 60 trials. In each trial, we generated 2,000 interferometric patterns at random axial positions ranging from ±500 nm and random cavity phase ranging from ± π rad with random lateral offsets of ±250 nm from the center within a sub-region size of 16 pixels. The photon count for each emission pattern was set to 2,000 and the background count per pixel was set to 5. To further validate the robustness, we introduced a deviation between the input phase and the ground truth in the construction of the 4Pi-BRAINSPOT model. Subsequently, we employed the model with these deviations to estimate the original data, setting the deviations at 0, π/4, π/2, and 3π/4 (30 trials for each deviation).

In conducting simulations of objective misalignment estimation, the same aberration configuration was used to generate 60 trials. In each trial, we generated 2,000 interferometric patterns at random axial positions ranging from ±500 nm and random objective misalignment ranging from ±160 nm laterally and ±50 nm axially (corresponding to contravariant tilt and defocus aberrations of ±1 λ/2π) with random lateral offsets of ±250 nm from the center within a sub-region size of 16 pixels. The photon count for each emission pattern was set to 2,000 and the background count per pixel was set to 5.

In conducting simulations of 4Pi channel-specific localization, we set the amplitude of vertical astigmatism to -2 and +2 λ/2π for the upper and lower pupils. We generated 1,000 interferometric emission patterns at 11 axial positions ranging from ±500 nm with a step size of 100 nm within a sub-region size of 16 pixels. The photon count for each emission pattern was set to 2,000 and the background count per pixel was set to 30. To simulate system imperfection, the simulations were sequentially rotated by 15 degrees using an affine transformation and incorporated Poisson noise processing as well as pixel-dependent Gaussian noise in sCMOS camera.

**Configuration of 4Pi-BRAINSPOT algorithm**

**3.1. Workflow of 4Pi *in-situ* PSF retrieval**

The 4Pi *in-situ* PSF retrieval algorithm in the 4Pi-BRAINSPOT approach, inspired by our previously published *in-situ* PSF retrieval method [15], captures interferometric distortions present in observed interferometric patterns. This algorithm constructs the 4Pi-PSF library by cropping individual interferometric emission patterns from acquired 4-channel single-molecule blinking images (Method 3.2) and assigns acquired patterns to their corresponding positions accurately in the absence of latent 3D positions (Method 3.3). These are fed into coherent 4Pi phase retrieval algorithm, retrieving the coherent pupil functions (Figure S3; Method 3.4). Through subsequent iterations, the coherent pupil functions were refined until stable coherent pupil functions were achieved. For the dynamic model update, we estimated and accommodated time-dependent interferometric aberrations according to objective misalignment in XYZ direction and cavity phase variation for each time window (Method 3.10&11). These coherent pupil functions were then used to construct a dynamic *in-situ* 4Pi-PSF model representing interferometric aberration information.

#### 3.2. Alignment of interferometric channels and construction of 4Pi-PSF library from single-molecule datasets

To ensure consistency across the dataset in 4-channel interferometric pattern processing, the raw 4-channel blinking datasets underwent an initial alignment prior to cropping emission patterns. This alignment involves using the maximum-intensity projection of 2,000 frames for each channel to calculate translation, scale, shear, and rotation operations between channel $p1$ and the other channels. These operations are estimated using affine transformation calculation via the 'imregtform' function in MATLAB. The resulting affine matrices are applied to align the other channels to channel $p1$ using the 'imwarp' function in MATLAB.

In cropping single-molecule patterns, the aligned datasets are superposed to relate the total acquired photon number across the FOV under incoherent conditions. To ensure each cropped pattern corresponding to individual single molecules, we follow a published configuration to check that maximum intensities exceed an initial intensity threshold $I_{\mathrm{init}}$, the center distance between any two nearby subregions is larger than a distance threshold $d_{\mathrm{thresh}}$, and the remaining intensities are higher than a segmentation threshold $I_{\mathrm{seg}}$ [15]. The coordinates of candidate single-molecule patterns are then used to crop interferometric emission patterns from the 4-channel dataset to construct the 4Pi-PSF library.

#### 3.3. Axial-position assignment and analysis of coherent interferometric patterns

The axial-position assignment began by generating reference 4Pi-PSF patterns at a series of Z positions. In the first iteration, the pupil functions of $h_U$ and $h_L$ are initiated with hyperparameters used in 4Pi-PSF simulation, including the aberration of vertical astigmatism set to ±1.5 λ/2π for simulation and ±1.2 λ/2π for experiment as prior knowledge. These were used to generate 4Pi-PSF patterns at various axial positions ranging from ±1 µm, with a step size of 50-100 nm. In subsequent iterations, the pupil functions $h'_U$ and $h'_L$ were updated based on the results from the previous iteration.

The interferometric patterns in 4Pi-PSF library were classified into distinct groups based on their similarity to the reference PSFs. The interferometric similarity score $S_{ij}$ was calculated between pair $i$ of reference 4Pi-PSF patterns $\mu_i$ and pair $j$ of acquired interferometric patterns $D_j$ across different channels by normalized cross-correlation (NCC) calculation, given by

$$S_{[illegible]} = \frac{1}{[illegible]} \sum NCC(\mu_{[illegible]}, D_{[illegible]}) \quad (18)$$

$$, where \ \ NCC(A,B) = \frac{1}{N}\sum_{n}\frac{1}{\sigma_A\sigma_B}A_nB_n$$

where NCC denotes the normalized cross-correlation operator, $n$ is value for pixel index $n$, $N$ is the total pixel number of $n$, $\sigma_A$ and $\sigma_B$ are standard deviations of $A$ and $B$. The interferometric patterns were assigned to a group with the highest similarity score. To ensure a robust analysis in following steps, we excluded interferometric patterns with a similarity score below $S_{min}$ of 0.5-0.6 and groups with a pair number below $N_g$ of 5-30. The interferometric patterns within each group were subjected to an alignment process and averaged to form a single 4Pi-PSF observation at a certain Z position, mitigating the noise effect. 2D cross-correlation analysis is operated to calculate the subpixel shift from the 4-channel reference 4Pi-PSF patterns, respectively, utilizing Fourier interpolation with a 10-fold up-sampling. Patterns of each channel are then aligned and normalized using z-score normalization before averaging operation.

#### 3.4. Principle of coherent 4pi phase retrieval

Coherent 4Pi phase retrieval algorithm was devised to retrieve hidden interferometric information from acquired intensity profiles of interferometric emission patterns. Our methodology employing Gerchberg–Saxton algorithm estimates coherent pupil functions of conjugate objectives from coherent interferometric patterns at a series of Z positions, following the relationships in Eq. 12&13 (Figure S3). To extract coherent patterns from partially coherent patterns, we employed prior knowledge of the coherence factor $a$ and the incoherent emission patterns, computed by either averaging the 4-channel patterns or simulating from initial pupil functions of upper and lower objectives.

The algorithm begins by initiating pupil functions of $h_U$ and $h_L$ with hyperparameters used in 4Pi-PSF simulation. The magnitude $A_m$ and phase $\Phi_m$ of coherent 4Pi-PSF amplitude $\mathrm{H}_m$ are calculated using Eq. 9-11. The amplitude $H'_m$ are updated by substituting magnitude $A_m$ with the acquired magnitudes of coherent interferometric patterns at corresponding z positions as

$$H'_m(x,y,z) = \sqrt{M_m(x,y,z)} \cdot e^{i\Phi_m(x,y,z)} \tag{19}$$

where $M_m(x,y,z)$ is the measured intensity profiles of coherent interferometric pattern at $z$ position. The interferometric wavefields $h'_m$ are updated by inverse Fourier transform of $H'_m$ as

$$h'_m(k_x,k_y,z) = \mathcal{F}^{-1}[H'_m(x,y,z)] \tag{20}$$

The coherent pupil functions $h'_U$ and $h'_L$ is then estimated from interferometric wavefields by solving the least-square minimization problem following the relationship in Eq. 9 as

$$\hat{X} = \arg\min\|Y - AX\|^2 \tag{21}$$

$$\text{where } Y = \begin{bmatrix} h'_{s1}(z_1) \\ h'_{s2}(z_1) \\ h'_{p1}(z_1) \\ h'_{p2}(z_1) \\ h'_{s1}(z_2) \\ \vdots \\ h'_{p2}(z_n) \end{bmatrix}, \ A = \begin{bmatrix} e^{i\pi}D(-z_1) & I_t e^{i(\varphi_{sp}+\varphi_0)}D(z_1) \\ D(-z_1) & I_t e^{i(\varphi_{sp}+\varphi_0)}D(z_1) \\ D(-z_1) & I_t e^{i(\pi+\varphi_0)}D(z_1) \\ D(-z_1) & I_t e^{i\varphi_0}D(z_1) \\ e^{i\pi}D(-z_2) & I_t e^{i(\varphi_{sp}+\varphi_0)}D(z_2) \\ \vdots & \vdots \\ D(-z_n) & I_t e^{i\varphi_0}D(z_n) \end{bmatrix}, \ \text{and } X = \begin{bmatrix} h'_U \\ h'_L \end{bmatrix}$$

The estimation of this overdetermined system can be derived by Moore–Penrose pseudo-inverse as

$$\hat{X} = (AA^T)^{-1}AY = \begin{bmatrix} h'_U \\ h'_L \end{bmatrix} \tag{22}$$

The updated coherent pupil functions $h'_U$ and $h'_L$ are used for the next iteration. After 60 iterations, the refined coherent pupil functions are regarded as the retrieved coherent pupil functions of the acquired interferometric emission patterns.

#### 3.5. Objective-misaligned interferometric aberration estimation

In the 4Pi microscopy system, objective drift in the XYZ directions results in interferometric aberrations, which can be categorized into covariant and contravariant terms [18]. These terms are orthogonal to each other, that any drift in the 4Pi system can be represented by a unique combination of them (Figure S8). In our study, common lateral drifts and opposing axial drifts were defined as covariant terms, while the opposite configuration was defined as contravariant terms. The impact of covariant drift was manifested as a translation of the 4Pi-PSF without inducing interferometric aberrations, whereas contravariant drift modified the interferometric patterns, thereby causing time-dependent discrepancies in the 4Pi-PSF. To deal with these independent impacts, we implemented a dynamic 4Pi-PSF model update to address time-dependent changes in the diffraction patterns and utilize a drift correction algorithm (Method 3.10) to correct time-dependent translations.

To address contravariant objective misalignment in the *in-situ* 4Pi-PSF model, the 4Pi blinking dataset is segmented into several time windows, each with a duration of 40 seconds. We assumed that within each window, the interferometric patterns exhibit consistent aberrations. For each time window, a 4Pi-PSF library was constructed following Method 3.2. Subsequently, a series of reference 4Pi-PSF patterns are generated based on devised coherent pupil functions with additional contravariant misaligned aberrations ranging from ±1 λ/2π (Method 2.6). To circumvent the phase wrapping issue, these misaligned interferometric aberrations were calculated directly in the phase term and integrated into the coherence pupil functions. Each interferometric pattern pair was assessed the similarity between all reference 4Pi-PSF models using Eq. 18, and assigned with a misalignment coefficient corresponding to the model exhibiting the highest similarity. We determined the overall coefficients of respective time windows by applying Gaussian fitting to pinpoint the peak of the distribution. The procedure was iterated across all dimensions to refine the coherent pupil functions, reducing discrepancies between the captured patterns and the *in-situ* 4Pi-PSF model. To ensure a robust analysis, we excluded interferometric patterns with a similarity score below $S_{min}$ of 0.5-0.6 and axial positions exceeding ±500 nm.

#### 3.6. Estimation of cavity-phase-induced interferometric aberration

The cavity phase characterizes the difference in optical path length from the common focal plane of the two objectives. To address time-dependent cavity phase variation, a dynamic *in-situ* 4Pi-PSF model update was incorporated across different time windows. We assumed that the interferometric patterns exhibit the same interferometric aberrations within each time window. For each time window, a 4Pi-PSF library was constructed following Method 3.2. Afterward, a series of reference 4Pi-PSF patterns were generated based on devised coherent pupil functions with additional cavity phase variation ranging from ±π rad (Method 2.6). Each interferometric pattern pair was assessed the similarity between all reference 4Pi-PSF models using Eq. 18, and assigned with a cavity phase corresponding to the model exhibiting the highest similarity. To enhance the accuracy of our analysis, particularly in addressing the phase wrapping issue observed around the boundary conditions of ±π rad, we employed periodic-padding Gaussian fitting. This method began with expanding the distribution of the cavity phase by shifting the phases ±2π rad from their original estimates, allowing a comprehensive representation of scores across cyclic cavity phases. We determined the overall coefficients of respective time windows by iteratively applying Gaussian fitting within a range of $\phi_{i-1} \pm \pi$ rad, where $\phi_{i-1}$ was estimated cavity phase in the previous iteration, until stable value was achieved. For the first iteration, $\phi_0$ was determined by mean of original estimates. To ensure a robust analysis, we excluded interferometric patterns with a similarity score below $S_{min}$ of 0.5-0.6 and axial positions exceeding ±500 nm.

#### 3.7. Principle of 3D channel-specific single-molecule localization

To preserve the statistical properties of the raw camera characterization, we generated channel-specific 4Pi-PSF models for each detected channel to enhance single-molecule localization accuracy and minimize potential imaging artifacts. The process began with mapping coordinates in reference channel $p1$ into the other channels using calculated affine matrices in Method 2.10. For non-integer coordinates resulting from the affine transformation, the transformed coordinates were rounded down to the nearest integer. The round-off center coordinate $(X'_m, Y'_m)$ for channel $m$, which was transformed from the center coordinate $(X_{p1}, Y_{p1})$ of the reference channel $p1$ using the affine transformation matrix $\begin{bmatrix} a & b & 0 \\ c & d & 0 \\ e & f & 1 \end{bmatrix}_m$, can be expressed as

$$(X'_m, Y'_m, 1) = \lfloor (X_m, Y_m, 1) \rfloor = \left\lfloor (X_{p1}, Y_{p1}, 1) \begin{bmatrix} a & b & 0 \\ c & d & 0 \\ e & f & 1 \end{bmatrix}_m \right\rfloor$$

$$\begin{cases} X'_m = \lfloor a_m X_{p1} + c_m Y_{p1} + e_m \rfloor = X_m - \Delta X_m \\ Y'_m = \lfloor b_m X_{p1} + d_m Y_{p1} + f_m \rfloor = Y_m - \Delta Y_m \end{cases} \quad (23)$$

where ⌊ ⌋ denotes the floor operator, $(X_m, Y_m)$ is the exact transformed coordinate, subscript $m$ is value for channel $m$, $(a, b, c, d)$ represent the scaling, shearing, and rotation operations, $(e, f)$ represents the translation operation, and $\Delta X$ and $\Delta Y$ are round-off errors in $X$ and $Y$ direction. The coordinate relationship between the center of subregions of channel $p1$ & $m$ can then be described as

$$(X'_m + x_m, Y'_m + y_m, 1) = (X_{p1} + x_{p1}, Y_{p1} + y_{p1}, 1) \begin{bmatrix} a & b & 0 \\ c & d & 0 \\ e & f & 1 \end{bmatrix}_m$$

$$\begin{cases} X'_m + x_m = a_m X_{p1} + c_m Y_{p1} + e_m + a_m x_{p1} + c_m y_{p1} \\ Y'_m + y_m = b_m X_{p1} + d_m Y_{p1} + f_m + b_m x_{p1} + d_m y_{p1} \end{cases} \quad (24)$$

where $(x_m, y_m)$ are the position of single molecules in the cropped subregions of channel $m$. By substituting Eq. 23 into Eq. 24, the relationship between $(x_{p1}, y_{p1})$ and $(x_m, y_m)$ can be expressed as

$$\begin{aligned} x_m &= a_m x_{p1} + c_m y_{p1} + \Delta X_m \\ y_m &= b_m x_{p1} + d_m y_{p1} + \Delta Y_m \end{aligned} \quad (25)$$

This analysis demonstrates that the cropped subregions, using round-off coordinates, share the same shape information, such as scale, shear, and rotation, as the original affine transformation. However, due to the rounding process, there may be translation discrepancies in the mapping from channel $p1$ to other channels. To compensate for these discrepancies, we applied a round-off calibration of ∆X and ∆Y for each channel, ensuring correct characterization of the sCMOS camera was applied. Characterization maps of the camera were also cropped for each channel according to the round-off subregion coordinates.

After mapping the coordinates, we generated 4-channel *in-situ* 4Pi-PSF patterns at a center position of $(x_{p1}, y_{p1}, z)$ using the coherent pupil functions retrieved from the *in-situ* 4Pi-PSF modeling process (Method 2.9). These patterns represent the 4Pi-PSF observations at the $z$ position with channel-specific phase delays near 0, π/2, π, and 3π/2. We then 2D transformed these *in-situ* 4Pi-PSF patterns $\mu_{p1}(x_{p1}, y_{p1}, z)$ into corresponding patterns $\mu_m(x_m, y_m, z)$ for channel $m$ to ensure consistency across all channels. The transformation with affine matrix $T_m$ calculated from Eq. 24&25 can be expressed as

$$\mu_m = Warp_{T_m}(\mu), \;\; T_m = \begin{bmatrix} a & b & 0 \\ c & d & 0 \\ \Delta X & \Delta Y & 1 \end{bmatrix}_m \quad (26)$$

where $Warp_{T_m}$ denotes geometric transformation operator using the 'imwarp' function in MATLAB with affine matrix $T_m$. This transformation accounted for the calibrated translation discrepancies due to the rounding process. As a result, transformed patterns exhibited the channel-specific *in-situ* interferometric fringe at the exact corresponding coordinates. This ensured that the presence of Poisson noise and pixel-dependent readout noise in the sCMOS camera was accurately represented for each channel.

To conduct single-molecule localization, we assumed the photon numbers are evenly distributed across the four detected channels to reduce axial localization uncertainty. We utilized a maximum likelihood estimator (MLE) with a likelihood function $L(\theta|D)$ integrating channel-specific 4Pi-PSF models and accounting for the presence of Poisson noise and pixel-dependent readout noise in the sCMOS camera, which can be described as:

$$L(\theta|D) = \prod_{m}\prod_{q}\frac{(\mu'_{m,q}+\gamma_{m,q})^{(D_{m,q}+\gamma_{m,q})}e^{-(\mu'_{m,q}+\gamma_{m,q})}}{\Gamma(D_{m,q}+\gamma_{m,q}+1)}, \theta \in (x,y,z,I,bg) \qquad (27)$$

where $\theta$ is the parameter set including 3D single-molecule center position $(x,y,z)$, photon number $I$, and background level $bg$, $D$ is the acquired photon value of cropped interferometric pattern, $\mu'$ is channel-specific 4Pi-PSF patterns, $\gamma = \frac{\sigma^2_{m,q}}{g^2_{m,q}}$ is the noise character of the sCMOS camera, and subscript $m,q$ is the value of pixel $q$ for channel $m$. The parameters $\theta$ can be estimated using the modified Levenberg-Marquardt method by solving the minimization problem of negative log-likelihood function

$$\hat{\theta} = \arg\min \ell(\theta) \qquad (28)$$

$$\text{where } \ell(\theta) = -\ln\big(L(\theta|D)\big) = \sum_{m}\sum_{q}\mu'_{m,q} - \big(D_{m,q}+\gamma_{m,q}\big)\ln\left(\mu'_{m,q}+\gamma_{m,q}\right) + C_{m,q}$$

where $C_{m,q}$ is constant of pixel $q$ for channel $m$ independent to $\theta$. The estimation of this minimization problem can be iteratively estimated by

$$\theta_{n+1} = \theta_n - \frac{f}{f'(1+\beta)}$$

$$\text{where}\begin{cases} f = -\dfrac{\partial \ln\big(L(\theta|D)\big)}{\partial\theta} = \displaystyle\sum_{m}\sum_{q}\left(1-\frac{D_{m,q}+\gamma_{m,q}}{\mu'_{m,q}+\gamma_{m,q}}\right)\frac{\partial\mu'_{m,q}}{\partial\theta} \\ f' = \dfrac{\partial f}{\partial\theta} = \displaystyle\sum_{m}\sum_{q}\frac{D_{m,q}+\gamma_{m,q}}{\left(\mu'_{m,q}+\gamma_{m,q}\right)^2}\left(\frac{\partial\mu'_{m,q}}{\partial\theta}\right)^2 + \left(1-\frac{D_{m,q}+\gamma_{m,q}}{\mu'_{m,q}+\gamma_{m,q}}\right)\frac{\partial^2\mu'_{m,q}}{\partial\theta^2} \end{cases} \qquad (29)$$

$f$ and $f'$ are 1st- and 2nd-order derivatives of negative log-likelihood function, and $\beta$ is a damping factor related to convergence speed. Here, we set second derivative of $\frac{\partial^2\mu'_{m,q}}{\partial\theta^2}$ and $\beta$ to be 0.

To evaluate the data-model similarity, the log-likelihood ratio (LLR) was calculated between acquired interferometric emission patterns and corresponding channel-specific interferometric patterns in Eq. 27 as

$$\text{LLR} = -2\ln\left(\frac{L(\mu'|D)}{\text{[illegible]}}\right) \qquad (30)$$

$$= 2\sum_{m}\sum_{q}\mu'_{m,q} - D_{m,q} - (D_{m,q} + \gamma_{m,q})\left(\ln(\mu'_{m,q} + \gamma_{m,q}) - \ln(D_{m,q} + \gamma_{m,q})\right)$$

#### 3.8. Precision evaluation using Cramér-Rao lower bound calculation

To evaluate the quality of acquired interferometric pattern, we used published configuration [19] to calculate the Fisher information matrix $\mathrm{F}(\theta_j)$ for estimated parameters $\theta_j$ under unbiased condition from the likelihood function in Eq. 27, which can be described as

$$\mathrm{F}(\theta_j) = \sum_{m}\sum_{q}\frac{1}{\mu'_{m,q} + \gamma_{m,q}}\left(\frac{\partial\mu'_{m,q}}{\partial\theta_j}\right)^2 \tag{31}$$

with Cramér-Rao lower bound (CRLB) precision $\sigma_\theta$ as

$$\sigma_\theta^2 \geq \frac{1}{\mathrm{F}(\theta_j)} \tag{32}$$

To calculate the Fisher information for channel-specific parameters, such as the single-molecule center position $(x_m, y_m)$ across all channels, we used the calibrated transformation relationship from Eq. 26 when calculating the dependent derivatives in the Fisher information. For example, considering the Fisher information $\mathrm{F_m}(x_{p1})$ for channel $m$, the 1st-order derivative can be expressed as

$$\frac{\partial\mu'_{m,q}}{\partial x_{p1}} = \frac{\partial\mu'_{m,q}}{\partial x_m}\frac{\partial x_m}{\partial x_{p1}} + \frac{\partial\mu'_{\mathrm{m},q}}{\partial y_m}\frac{\partial y_m}{\partial x_{p1}} = a_m\frac{\partial\mu'_{m,q}}{\partial x_m} + \mathrm{b_m}\frac{\partial\mu'_{\mathrm{m},q}}{\partial y_m} \tag{33}$$

This leads to the following expression for the Fisher information for channel $m$ as

$$\mathrm{F_m}(x_{p1}) = \sum_{q}\frac{1}{\mu'_{m,q} + \gamma_{m,q}}\left(a_m\frac{\partial\mu'_{m,q}}{\partial x_m} + \mathrm{b_m}\frac{\partial\mu'_{\mathrm{m},q}}{\partial y_m}\right)^2 \tag{34}$$

Following the same process, the Fisher information $\mathrm{F_m}(y_{p1})$ for channel $m$ can be expressed as

$$\mathrm{F_m}(y_{p1}) = \sum_{q}\frac{1}{\mu'_{m,q} + \gamma_{m,q}}\left(c_m\frac{\partial\mu'_{m,q}}{\partial x_m} + d_m\frac{\partial\mu'_{m,q}}{\partial y_m}\right)^2 \tag{35}$$

#### 3.9. Addressing ambiguities in 4Pi *in-situ* PSF retrieval

Ambiguity in 4Pi-SMSN appears when multiple sets of variables lead to the same observation of interferometric patterns. Here, we examined the ambiguity issues because of the symmetric nature of the 4Pi approach and how our methodology addressed the ambiguity issues to ensure clarity in analyzing the interferometric patterns.

The first ambiguity in 4Pi-SMSN arises from the axial indistinguishability when the 4Pi-PSF exhibits axial symmetry because of coupling between axial position and symmetric interferometric aberrations. For instance, the interferometric patterns of a single molecule at z = 600 nm, with vertical astigmatisms of ±1 λ/2π in the upper and lower pupil functions, appear identical to the patterns observed at z = −600 nm with vertical astigmatisms of ∓1 λ/2π (Figure S8A). This ambiguity poses challenges to axial-position assignment process in the 4Pi phase retrieval algorithm, as interferometric patterns cannot be uniquely assigned to specific axial positions without additional information about coherent pupil functions or ground-truth single-molecule axial positions. To resolve this issue, we introduced vertical astigmatism with specific amplitudes as prior knowledge, breaking the indistinguishability between axial position and symmetric interferometric aberrations.

The second ambiguity arises from the distinguishability of coherent pupil functions, when interferometric aberrations contribute equally to both pupil functions. For instance, the interferometric patterns with vertical coma aberrations of -0.5 and 1.5 λ/2π in the upper and lower pupil functions, respectively, appear identical to the patterns with coma aberrations of -1.5 and 0.5 λ/2π (Figure S8B). This ambiguity poses challenges in the coherent pupil functions update process in the coherent 4Pi phase retrieval algorithm, as interferometric wavefields cannot be uniquely decomposed into specific coherent pupil functions without information about ground-truth pupil functions. Although this ambiguity cannot be broken up under 4Pi-BRAINSPOT framework, the distinguishability of coherent pupil functions does not influence the uniqueness in the representation of the 4Pi-PSFs. To prevent this ambiguity in the localization process and other analyses, we exclusively focused on interferometric patterns when interpolating the 4Pi-PSF model.

Another ambiguity arises from the distinguishability of coherent pupil functions, particularly when translating interferometric aberrations induced by common-term lateral objective drifts and opposite-term axial objective drifts. For instance, the interferometric patterns with horizontal tilt aberrations of 0 and 4 λ/2π in the upper and lower pupil functions, respectively, appear identical to the patterns with horizontal tilt aberrations of -2 and 2 λ/2π once we ignore the translation (Figure S8C&D). This ambiguity poses challenges in estimating objective-misaligned interferometric aberrations in the dynamic 4Pi-PSF model update process, as translating interferometric aberrations cannot be uniquely determined without information about ground-truth coherent pupil functions or ground-truth single-molecule positions. Although this ambiguity cannot be resolved within the 4Pi-BRAINSPOT framework, the translating interferometric aberrations don't influence the uniqueness in the representation of the 4Pi-PSFs. To prevent this ambiguity, we excluded translating interferometric aberrations in the dynamic 4Pi-PSF model update process and compensated for its influence through drift correction at the end of the localization process. After removing the translation ambiguity, we can utilize the unique amplitude of objective misalignment, including x tilt, y tilt and defocus, to characterize the overall movement of the objectives (Figure S4A, Method 3.5).

#### 3.10. Drift correction in 4Pi-BRAINSPOT reconstruction and evaluation of localization reliability

To ensure the reliability of the reconstructed images from 4Pi single-molecule localization, we applied filtering criteria based on the CRLB, LLR, axial range of localization, and photon counts. We excluded localizations of interferometric patterns with a CRLB estimation above 10 nm for all three dimensions, an LLR below 2400, axial positions exceeding ±600 nm, and photon numbers below 1500 to ensure that only high-quality localizations contribute to the reconstructed image.

Drift correction was carried out using a redundant cross-correlation configuration [15]. We divided the localizations into *n* segments, with each segment containing between 1,000 and 2,000 frames, and rendered them as *n* 3D volumes. 3D cross-correlation analysis was operated to calculate the subpixel shift between any two segments utilizing Fourier interpolation with a 10-fold up-sampling, resulting in total $\frac{1}{2}n(n+1)$ shift estimations. We determined the $n$-1 unknown 3D shift between adjacent segments from the overdetermined estimations and performed 3D drift correction to each segment. To further enhance the alignment accuracy in 3D reconstruction, additional drift correction was performed along the xz, yz, and xy directions, respectively.

#### 3.11. Imaging rendering of 4Pi-BRAINSPOT reconstruction

In imaging rendering of 2D images using MATLAB, the localizations were divided into 64 segments based on their axial positions. Depending on rendering transverse or vertical cross section, lateral or axial localization coordinates in each segment were mapped onto 2D images with a pixel size of 2-3 nm and pseudo-colored according to the axial position of the segment. In cases where multiple localizations were located at the same pixel, their values were accumulated. To blur the images, we applied a 2D Gaussian blur with 0.7-1 RMS width of overall CRLB precision. These blurred images were then merged to form a composite image. To enhance image contrast, pixel intensities were linearly scaled from the original range [0, $L_{high}$] to [0, 255], where $L_{high}$ is a determined upper limit for imaging rendering. Lastly, we further stretched the image contrast to saturate the top 1-5 % of pixels with the highest intensity values, minimizing contrast variation caused by exceptionally bright pixels.

In the 3D reconstruction process using Blender, we represented each localization as a particle positioned at its corresponding coordinates. These particles were pseudo-colored based on their axial positions to provide depth information. To visually represent the uncertainty associated with each localization, we rendered each particle as a two-layered sphere. The inner sphere had a diameter equal to the CRLB precision and was rendered with high opacity, while the outer sphere, with a diameter twice that of the inner sphere, was rendered with low opacity. To enhance the overall image contrast and reduce the impact of exceptionally bright pixels, we saturated the top 1-5 % of pixels with the highest intensity values.

### Quantification and statistical analysis

#### 4.1. Analysis pipeline of 3D structural morphology and circumference

To accurately measure the morphology along interested structures from single-molecule localizations, we devised a semi-auto analysis pipeline for spine quantification. This process began by mapping the localization coordinates onto a 2D image with a pixel size of 10-20 nm, allowing users to identify the general spatial locations of the structures of interest by marking along them. The algorithm then traced these marks as a smooth curve using the 'cubicspline' fitting method in MATLAB and generated segments along the curves with dimensions of 200-600 nm in width, 600-1000 nm in height, and 150 nm in length. To prevent structural discontinuities due to the inherent sparsity of 4Pi-SMSN, we introduced a 50-nm overlap between adjacent segments.

Within each segment, the algorithm delineated the major structure using the 'alphaShape' algorithm in MATLAB, generating complex boundaries by uniting multiple convex hulls with an alpha radius of 0.3-0.5 in the cross-section encompassing subsets of localizations. Localizations falling within the structural region were fitted with a 2D elliptical function to locate the structural center. To minimize the impact of random outliers in radius measurement, we measured the distances between the structural center and all localizations within each segment, generating histograms with a bin size of 4-8 nm, and identified the mode of distances by fitting the histogram with a Gaussian function. The mode of distances was regarded as the geometric mean of radius. The volume trace was generated by fitted circular areas calculated from the geometric mean of radius using the 'smoothingspline' fitting method in MATLAB. To quantify the distribution of localizations on a structural surface, we sliced the straightened structures along their longitudinal direction and flattened them into a 2D surface. The localizations were then mapped to their corresponding angular positions, generating a 2D circumference plot color-coded according to angular positions ranging from ±π.

#### 4.2. Analysis of PSF relevance and data visualization

To compare the similarity between two diffraction patterns, we calculated the similarity score $S_{3D}$ between two interferometric patterns $A$ and $B$ in three dimensions by Eq. 18 as

$$S_{3D} = \frac{1}{4Z} \sum_{z} \sum_{m} NCC\left(A_{m,z}, B_{m,z}\right) \tag{36}$$

where $A_{m,z}$ and $B_{m,z}$ are interferometric patterns at $z$ position for channel $m$., and $Z$ is total number of $z$. In the evaluation of localization uncertainty, we assessed the accuracy and precision of our system through fluorescent bead measurement. The accuracy and precision at each step were determined by calculating the mean absolute deviations and standard deviations, respectively, between the localized positions of the beads and their known ground truth positions.

In a box-chart visualization, the central line represented the median of the dataset and box represented upper and lower quartiles. The whiskers extended from the box to the furthest data points excluding outliers. Outliers were defined as values greater than $q_3 + 1.5 \times (q_3 - q_1)$ or less than $q_3 - 1.5 \times (q_3 - q_1)$. In a bar chart visualization, the bar heights represented the mean of the dataset, while the error bars indicated the standard deviation. For statistical significance, the p-values were calculated using 'ttest2' function in MATLAB, where the p-values smaller than 0.001 are denoted as $p<0.001$.

#### 4.3. Measurement of illumination profile in 4Pi-BRAINSPOT and calculation of signal-to-background ratio

The dye-on-coverslip sample was illuminated with a modest laser power of 30 W/cm$^2$ for imaging acquisition, after which system was calibrated employing single-molecule fluorophores on coverslips (Method 1.10). To measure the tilted angle of illumination from the lower objective, the illumination area was reduced to a diameter of 7 μm, preventing clipping by FOV. The dye samples were scanned within an axial range of ±25 μm and acquired using upper objective with a synchronized scan. The step size for this scan was 2 μm, and each position was recorded for one frame with an acquisition time of 500 ms. For each measured image, illuminated regions were identified by calculating the median intensity as a threshold. We marked pixels with intensities above this threshold and generated patterns to encompass the marked pixels. The patterns were fitted with a 2D elliptical function to determine their centers and diameters at different axial positions. The tilted angle of the illumination profile was subsequently calculated based on the center orientation of the illumination within an axial range of ±15 μm.

To estimate the Signal-to-Background Ratio (SBR), we simulated a 40x40x50 μm$^3$ sample volume, with a unit cube size of 10 nm, containing uniformly distributed fluorophores. We assumed the imaging plane was located at the center of this volume and that fluorescence emission was linearly proportional to the illumination intensity. In the case of epi-illumination, we simulated uniform illumination throughout the entire volume. For HILO illumination, we modeled a light sheet with a Gaussian intensity profile perpendicular to its propagation direction, with a $1/e^2$ width of 22.4 μm and intersecting the sample at an angle of 54° to the optical axis. During detection, the FOV was 20 μm, and fluorescence originating within an axial range of ±800 nm from the imaging plane was considered the signal. Fluorescence originating from regions outside this range was considered background. The detected emissions were calculated by aggregating contributions from different depths. For each depth, the fluorescent contribution was simulated by convolving the out-of-focus PSF pattern with the fluorescence map, which was determined as the product of the fluorophore density and the illumination profile. Consequently, the SBRs of 54°-HILO and 0°-Epi illumination were 8.0% and 3.2%, respectively, indicating a 2.5-fold improvement with a 60% reduction in background volume for a 50-μm tissue sample. Following the same simulation for an 80-μm-thick sample, it would be a 4.3-fold SBR improvement with a 77% background reduction.

#### 4.4. Measurement of intensity profile from 4Pi-SMSN dataset

To accurately measure the intensity profile of single-molecule localizations, we applied a histogram measurement to the localizations. This process began with mapping the localization coordinates onto a 2D image with a pixel size of 1.3 nm, enabling users to manually identify the structures of interest. The algorithm generated box segments along the line with a size of 4-6 nm and accumulated the number of localizations inside each segment, creating histogram profiles along the structures. To quantify the features, we fitted $n$ Gaussian functions to $n$ local maximum in the histogram, whose centers and FWHMs were regarded as the center positions and widths of features. To ensure robust analysis, we excluded features with localization counts below 2 and widths below CRLB-estimated resolution. This approach obtained a nanoscale measurement of the intensity profile by minimizing imaging artifacts caused by the blurred kernel in the rendering process. To compare the intensity profiles of the same structures using various localization methods, we initially aligned the reconstructions using a redundant cross-correlation configuration and drew lines across similar structures, minimizing differences in the profiles due to global shifts between the reconstructions.

#### 4.5. Quantification of cluster of single-molecule localization

To identify single-molecule clusters, we assumed that localizations originating from a single molecule follow the 3D Gaussian random process with standard deviations of the overall CRLB precisions. The single-molecule clusters were determined once subsets of localizations fell within three standard deviations with no localization between three and five standard deviations. Statistically, when there are 5 observations in a cluster, the hit probability that all localizations will fall within three standard deviations is 98.7%. Conversely, the miss probability that at least one localization will fall between three and five standard deviations is 1.3%. To evaluate the size of clusters, the identified clusters were aligned using their center positions and superposed as a single cluster. The size of the superposition was determined by fitting with 2D Gaussian functions to calculate FWHM. To validate the accuracy of cluster size measurements, we also evaluated mean and standard deviation across all cluster sizes. To ensure robust analysis, we excluded clusters with a localization number below 4 and a total photon number below 8000.

#### 4.6. Quantification of 4Pi-SMSN resolution

In this work, three different resolution quantifications were used. In the directional Fourier correlation (DFC) analysis, we segmented the localizations into 11x11 grids based on their lateral coordinates. Within each segment, the localizations were randomly divided into two statistically independent subsegments. The localization coordinates of each subsegment were mapped onto 3D images with a volume size of 3 nm. To prevent boundary issues, a 3D Tukey window function was applied to each 3D image. For each segment, the spatial frequency components along the certain direction of the two subsegments were extracted for correlation calculation. For a spatial frequency vector $\vec{q}$, the directional Fourier correlation was defined as

$$DFC(\vec{q}) = \frac{\sum_{i \in I} \widehat{f_{\iota,1}}(\vec{q}) \widehat{f_{\iota,2}}(\vec{q})^*}{\sqrt{\sum_{i \in I} \left| \widehat{f_{\iota,1}}(\vec{q}) \right|^2 \ \sum_{i \in I} \left| \widehat{f_{\iota,2}}(\vec{q}) \right|^2}} \tag{37}$$

where $\widehat{f_{\iota,1}}(\vec{q})$ and $\widehat{f_{\iota,2}}(\vec{q})$ were the spatial frequency components at frequency $\vec{q}$ in subsegments of segment $i$ and $i \in I$ encompasses all valid segments. The resulting DFC curves were then smoothed using the 'cubicspline' fitting method in MATLAB to reduce noise and facilitate accurate resolution determination. The DFC resolution for each segment was determined as the inverse of the spatial frequency at which the smoothed DFC curve reached the 1/7 criteria. To ensure robust analysis, segments were excluded with localization numbers below 200.

The Fourier shell correlation (FSC) and imaging decorrelation analysis were performed based on published configurations [20,21]. In the FSC analysis, we segmented the localizations into 11x11 grids based on their lateral coordinates. Within each segment, the localizations were randomly divided into two statistically independent subsegments. The localization coordinates of each subsegment were mapped onto 3D images with a volume size of 3 nm. To prevent boundary issues, a 3D Tukey window function was applied to each 3D image. The FSC calculation [20] for each segment was performed between the 3D Fourier transforms of the two subsegment images. This process involves computing the cross-correlation of corresponding shells in Fourier space and normalizing it by auto-correlation of each shell. The resulting FSC curves were then smoothed using the 'cubicspline' fitting method in MATLAB to reduce noise and facilitate accurate resolution determination. The FSC resolution for each segment was determined as the inverse of the spatial frequency at which the smoothed FSC curve reached the 1/7 criteria. To ensure robust analysis, segments were excluded with localization numbers below 200.

In the imaging decorrelation analysis [21], we also segmented the localizations into 11x11 grids based on their lateral coordinates. Within each segment, the localization coordinates were mapped onto monotonic 2D images with a pixel size of 3 nm and blurred using a 2D Gaussian blur with RMS width of overall CRLB precision. The decorrelation analysis for each segment involved computing the cross-correlation between the Fourier transform of the image and its binary circular mask-filtered version, under a series of Fourier low-pass and high-pass filtering. The imaging decorrelation resolution for each segment was determined by $\frac{2 \times Pixel\ size}{cut-off\ frequency}$. This cut-off frequency identifies the highest spatial frequency at which a local maximum occurs in the decorrelation curves, indicating the spatial frequency beyond which the signal is no longer preserved. To ensure robust analysis, segments were excluded with localization numbers below 200.

### Data and software availability

The 4Pi-BRAINSPOT toolbox, along with detailed instructions, is available as a supplemental ZIP file accompanying the submission. Updates to the toolbox will be continuously made available freely on the Github repository as of the date of publication. The software package features an easy-to-use user interface including all steps of 4Pi reconstruction workflow from channel alignment, 4Pi-PSF segmentation, 4Pi in situ PSF retrieval, dynamic model update, 4Pi localization (supporting pupil-based 3D localization with cubic spline GPU implementation), drift correction, volume alignment, to super-resolution image reconstruction. The data are available at Mendeley Data. Any additional information required to analyze the data is available from the lead contact on request.
